# Systematic Infectome–Phenome Profiling Reveals Cryptococcal Infection-Associated Proteins Driving Immune System Remodeling and Immunization Potential

**DOI:** 10.64898/2026.01.30.702846

**Authors:** Brianna Ball, Norris Chan, Hannah West, Arjun Sukumaran, Duncan Carruthers-Lay, Michael Woods, Jennifer Geddes-McAlister

## Abstract

Mortality and morbidity associated with invasive fungal infections continue to rise, driven by the emergence of new pathogens, increasing antifungal resistance, and expanding immunocompromised populations. Despite this growing threat, fungal biology lacks scalable, unbiased strategies to link infection-responsive fungal proteins to functional outcomes that drive virulence and immune modulation. Here, we present an integrated infectome–phenome discovery platform that combines high resolution mass spectrometry–based proteomics with systematic phenotypic profiling to globally interrogate *Cryptococcus neoformans*–macrophage interactions. This approach reveals coordinated host immune suppression alongside complementary fungal virulence programs and enables the unbiased prioritization of infection-associated fungal proteins. Systematic phenome fingerprinting of candidate mutant strains resolves two functional classes of putative therapeutic relevance: antifungal and antivirulence, with *in vitro* characterizations corroborated with a murine model of cryptococcosis. Prioritization of a conserved, previously uncharacterized virulence-associated protein, CipC, uncovers altered extracellular vesicle composition and enhanced antigenic properties. Immunization with CipC-derived vesicles elicits a robust and diversified host immune response, implicating fungal extracellular vesicles in immune remodeling and host priming. Together, these findings establish a broadly applicable framework for systematically identifying and functionally characterizing fungal drivers of infection for therapeutic and immunological target discovery with relevance across diverse human fungal pathogens.

## Introduction

Fungal infections represent a growing global health challenge, affecting millions of individuals each year and ranging from superficial disease to invasive, life-threatening infections. While fungi have long been considered opportunistic pathogens primarily affecting immunocompromised individuals, including those with co-infections, immunomodulatory therapies, and advanced age, recent trends indicate a concerning rise in infections among immunocompetent populations^1^. At the forefront of this challenge is *Cryptococcus neoformans,* a major invasive opportunistic fungal pathogen and the primary causative agent of cryptococcal meningitis^2,3^. It is responsible for ∼180,000 global annual deaths, and the mortality rate in resource-limited countries exceeds 70%^4,5^. Treatment of these invasive fungal infections encounters many challenges, including high host cytotoxicity from shared homology of drug targets, a limited antifungal arsenal, growing populations of at-risk individuals, and the emergence of resistant strains to clinically available antifungal drugs^6^. Therefore, it is imperative to elucidate novel antifungal targets and strengthen the antifungal pipeline.

*C. neoformans* primary infection initiates with pulmonary colonization, where fungal cells encounter the first line of host immune defense: resident alveolar macrophages. *C. neoformans* employs multiple sophisticated virulence traits, including the production of a polysaccharide capsule, generation of melanin pigment, and extracellular enzyme production, which combat host immune cells^7–10^. These fungal virulence factors enable evasion of phagocytosis and survival within the macrophage phagolysosome, whereas subsequent escape or transfer between macrophages may also occur, often without host cell death^11–13^. Extrapulmonary dissemination of the fungus occurs hematogenously, with eventual crossing of the blood-brain barrier (BBB) and invasion of the central nervous system (CNS) leading to meningitis or meningoencephalitis^14^. Throughout the infection stages, the interactions of cryptococcal cells with macrophages remain constant and are considered a primary determinant of infection outcome^15^. Despite the abundant information on the interplay between *C. neoformans* and macrophage, we have yet to comprehensively profile and understand critical fungal and macrophage effectors involved in infection.

Dual perspective proteome profiling (i.e., detection and quantification of host and pathogen proteomes during infection), is a powerful approach to understand how the host defends itself from invasion and how the pathogen overcomes these defenses^16,17^. Recent exploration for our research team and others, have applied this approach using mass spectrometry-based proteomics to identify fungal-activated immune responses^18^, propose biomarkers and diagnostic markers of cryptococcal infection^19^, define putative antifungal targets^20^, and explore protein-level drivers of antifungal resistance^21^. Beyond these global studies, a systematic approach to distinguish growth-vs. infection-associated cryptococcal proteins are limited to genetic deletion studies and assessments of *in vitro* and *in vivo* virulence^22,23^. While such studies are highly informative for the identification of individual putative novel virulence factors from the genetic perspective, systematic investigation of the proteins that are produced to support dynamic infection and survival of the pathogen within the host are lacking. This information supports the design of therapeutics that specifically target fungal proteins relevant during infection to disarm the pathogen and empower the host to clear the infection. Additionally, strategies to better characterize the role of infection-relevant fungal proteins for activation of the immune response and protection against disease can also be developed.

In this study, we present an integrated infectome–phenome discovery platform that combines mass spectrometry–based proteomics with systematic phenotypic profiling to globally interrogate *C. neoformans*–macrophage interactions. This approach uncovers coordinated host immune suppression alongside complementary fungal virulence programs, addressing a major gap in cryptococcal biology: the lack of scalable, unbiased strategies to link infection-responsive fungal proteins to functional outcomes. Herein, we used high resolution mass spectrometry-based proteomics to reveal the host and fungal counterparts influenced during infection. We unbiasedly observe cryptococcal-mediated host immune suppression and complementary fungal virulence-protein signatures. We prioritized 15 fungal infection-associated proteins measured as strictly macrophage-responsive and systematically phenotyped candidate mutant strains for *C. neoformans* classical virulence factors. This phenome fingerprinting defined two categories of putative fungal therapeutic candidates, including antifungal and antivirulence. Assessment of a prioritized candidate from each of the fungal therapeutic categories in a murine model of cryptococcosis revealed corroboration across murine survival to the *in vitro* phenome. Critically, our novel therapeutic target selection pipeline discovered an uncharacterized, conserved, and virulence-associated protein target, CipC (CNAG_03007) with altered extracellular vesicle (EV) signatures. Importantly, heightened production of antigenic properties within CipC vesicles did not protect the mice against cryptococcal infection; however, we observed induction of a robust immune response, supporting a new role in host immunization for diverse human fungal pathogens. Our findings define a new platform for the identification and characterization of fungal drivers of infection and methodically designates and evaluates prime novel therapeutic targets for improved strategies to treat and protect the host against fungal pathogens.

## Results

### Dual-perspective proteomic profiling prioritizes fungal infection-associated proteins

The outcome of cryptococcosis is dictated by fungal virulence traits leveraging and manipulating intrinsic macrophage capabilities, including fungal residence of the phagosome and resulting widespread circulation throughout the body^24^. To characterize host and fungal responses to infection, we performed quantitative proteomics profiling during *C. neoformans* infection of primary monocytic-derived macrophages (i.e., infectome) in comparison to the ‘*in vitro’* fungal proteome (i.e., fungal growth within enriched media). The *in vitro* fungal proteome and secretome identified 2767 and 18 exclusive proteins with 25 proteins shared between the cell pellet and extracellular environment (Fig. 1A); these proteins are produced by *C. neoformans* in growth media (in the absence of macrophages). Next, dual-perspective proteome profiling of the fungal-macrophage infection, capable of species-specific protein identification within a single mass spectrometry run, detected a distribution of 15% fungal (874) and 85% host (4952) proteins (Fig. 1B). The infectome profiling revealed strong reproducibility across biological replicates (Infected replicates = 95.6%; Uninfected replicates = 96.6%) (Fig. S1) and clear separation by principal component analysis (PCA) between infected and uninfected macrophages (component 1, 44.66%) with biological variability accounting for the second component of distinction (component 2, 16.99%) (Fig. 1C). A volcano plot of significantly different host proteins revealed 214 proteins with increased abundance within uninfected samples and 73 proteins with increased abundance in the infected samples (Fig 1D; Table S1).

**Fig. 1:**
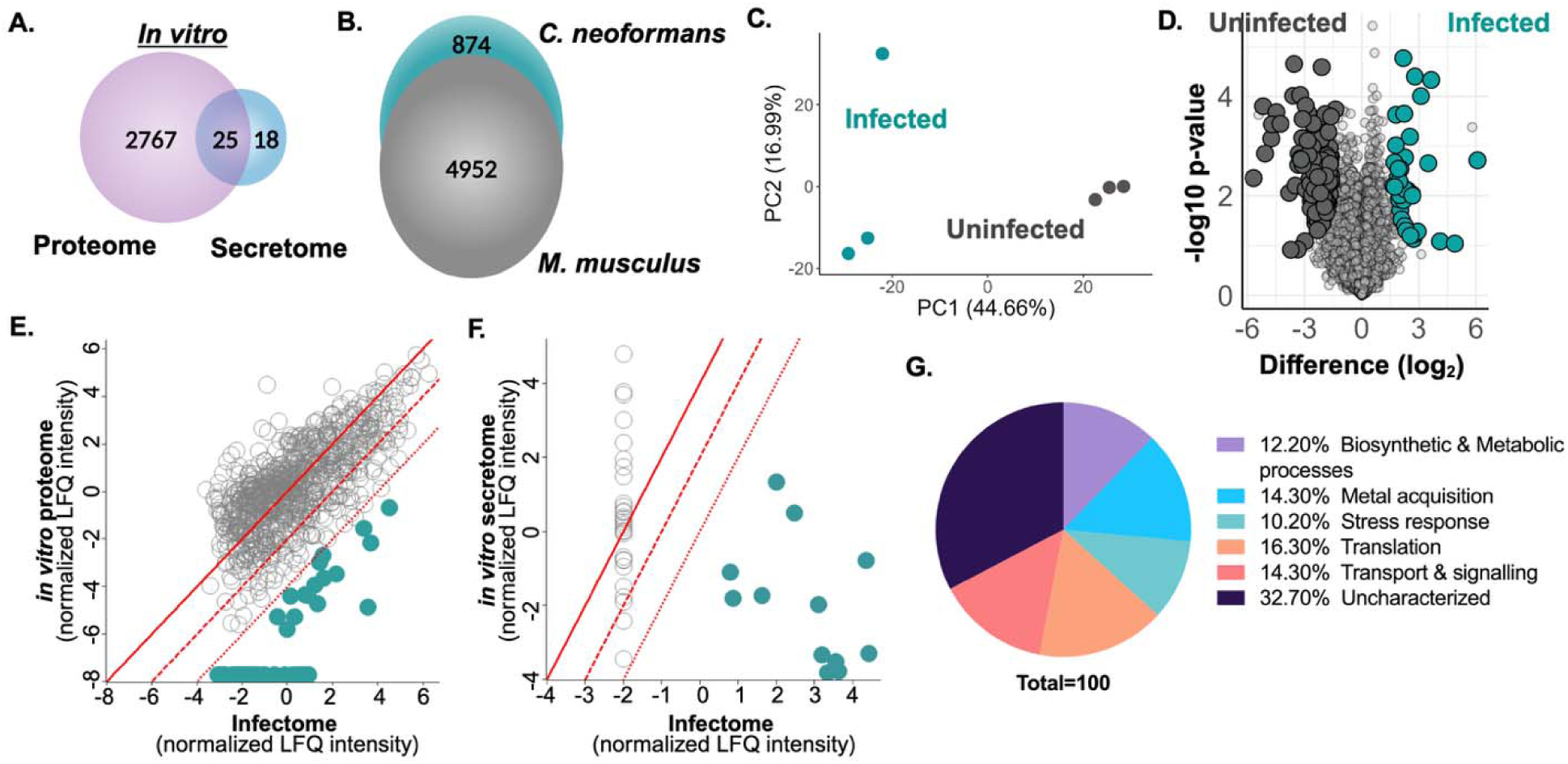
Dual-perspective proteome profiling detects and prioritizes infection-associated fungal proteins. **A**) Venn diagram of unique proteins identified in the *in vitro* cellular proteome (purple; 2767 proteins) and secretome (blue; 18 proteins) with 25 proteins commonly detected. **B)** Egg diagram of proteins identified from the host, *Mus musculus* (4,952 after valid value filtering), and pathogen, *Cryptococcus neoformans* (874 after valid value filtering) from infectome analysis. **C)** Principal component analysis of infectome experiment. **D)** Volcano plot for comparison of host proteins with significant differences in abundance between infected (teal) and uninfected (grey) conditions. Student’s *t* test, *P* < 0.05; FDR = 0.05; S_0_ = 1. **E)** Comparison of significantly different fungal proteins identified from the *in vitro* cellular proteome compared to the infectome. **F)** Comparison of significantly different fungal proteins identified from the *in vitro* secretome compared to the infectome. Red lines represent 0-, 4-, and 16-fold change in protein abundance. **G)** Pie chart based on GOBP for 53 fungal infection-associated protein (percentage of each category represented). Experiment completed in biological triplicate.

Given the study’s objective to identify and characterize fungal infection-associated proteins, we compared the *in vitro* cellular proteome and secretome of *C. neoformans* to the infectome. We filtered for fungal proteins with a ≥16-fold increase in abundance or proteins exclusively detected during macrophage co-culture to ensure prioritization of infection-relevant fungal proteins. We reported 41 fungal proteins meeting these criteria upon comparison to the cellular proteome (Fig. 1E) and 12 fungal proteins from the secretome (Fig. 1F; Table S2). GOBP assignment for the 53 (49 upon removal of proteins identified in both cellular proteome and secretome) infection-associated proteins defined distribution across biosynthetic and metabolic processes (e.g., dehydratase), metal acquisition (e.g., copper acquisition factor^25^), stress response (e.g., superoxide dismutase, heat shock 90 co-chaperone ^26–30)^, translation (e.g., initiation factor), transport (e.g., alternative oxidase), and uncharacterized (Fig. 1G). Next, we selected 17 candidate proteins for functional follow-up based on novelty, potential roles in virulence, and predicted secretion or membrane localization. To verify production of these fungal proteins as a result of macrophage presence and not media differences (e.g., YPD vs. DMEM), we confirmed increased abundance for 15 of the 17 selected fungal candidates (Fig. S2A, S2B). Overall, our comparison of *in vitro* vs. macrophage-infection fungal proteomes revealed prioritized infection-relevant targets for systematic investigation.

### Systematic phenotyping of novel infection-associated fungal candidates

To evaluate the roles of the prioritized fungal proteins on virulence, we generated and confirmed gene deletion strains for the 15 candidates. We performed *in vitro* systematic phenotypic profiling for *C. neoformans* classical virulence factors, including polysaccharide capsule production, melanization, thermotolerance, and release of lactate dehydrogenase upon macrophage infection. We identified 11 mutant strains with reduced capsule compared to WT, including *CNAG_05069*Δ and *CNAG_05997*Δ (associated with the electron transport chain, mitochondrial function, and capsule growth^31^) and *CNAG_03007*Δ (a CipC protein homolog associated with filamentation and putatively cell wall integrity in *Aspergillus fumigatus* ^32,33^)

(Fig. 2A; Fig. S3A, S3B). Defects in melanization were observed for 11 mutant strains, including candidates previously reported to display melanin defects, *sod1*Δ and *bim1*Δ^25,34^; melanin-association for the remaining nine candidates was not previously reported (Fig. 2B; Fig. S3C). Next, we evaluated fungal growth and thermotolerance in synthetic minimal media (YNB) at 30°C and 37 °C, respectively. We observed a severe thermotolerance defect for *CNAG_05997*Δ and moderate thermotolerance defects for seven additional mutant strains (Fig. 2C; Fig. S3E, S3F). To further assess the infection-associated candidates’ role in virulence, we quantified lactate dehydrogenase (LDH) release by macrophages following co-culture with the mutant fungal strains. We observed 15 mutant strains with reduced host cell cytotoxicity compared to WT (Fig. 2D; Fig. S3D).

**Fig. 2:**
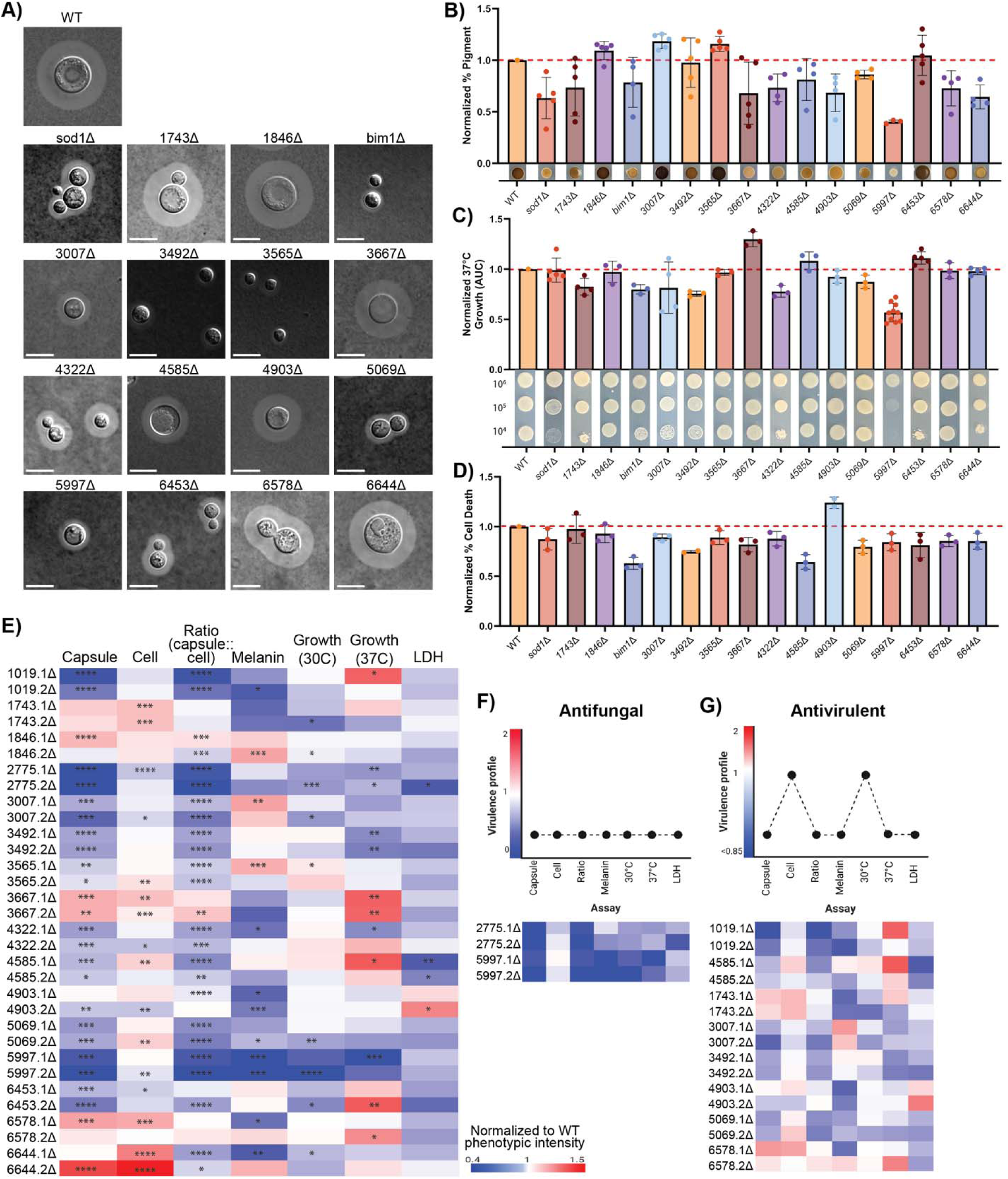
Systematic phenome profiling of *C. neoformans* infection-associated candidates. **A)** Capsule production of infection-associated mutant strains. Strains grown in iron-limited capsule inducing media to mid-log phase (∼24 h), visualized by India Ink staining with differential interference (DIC) microscopy. Representative images of mutant strains shown. Scale bar = 4.6 µm. A minimum of 50 cells were measured for capsule and cell size assays across three biological replicates. **B)** Normalized percentage of melanin (pigment) produced by mutant strains. Experiment performed in biological duplicate and technical triplicate. Representative melanized colonies shown below figure. **C)** Normalized 37 °C growth profiles obtained from area under the curve (AUC) measurements of mutant strain growth curves in YNB. Representative spot dilution plates shown below figure. Experiment performed in biological triplicate or quadruplicate and technical duplicate. **D)** Normalized percentage of host cell death by quantification of lactate dehydrogenase (LDH) release from BALB/c macrophages at 27 hpi following co-culture with infection-associated mutant fungal strains. Mutant strains opsonized by monoclonal 18b7 antibody prior to co-culture. Experiment completed in biological triplicate and technical duplicate or triplicate. **E)** Phenome heat map of infection-associated mutant fungal strains. Phenome scores generated by normalization against WT based on quantitative or semi-quantitative measurements. **F)** Categorization of candidates as ‘antifungal’ based on defect across all evaluated criteria. **G)** Categorization of candidates as ‘antivirulent’ based on defect across classical virulence factor criteria. Red and blue schemes represent enhanced and reduced phenotype scores, respectively. Significance determined by Students *t* test: *P* < 0.05, *; *P* ≤ 0.01, **; *P* ≤ 0.005, ***; *P* ≤ 0.0001, ****. Phenotypic assays performed with two independent mutant strains; figure generated with representative single independent mutant strain.

### Integration of phenome fingerprinting across virulence factors defines antifungal and antivirulence categories

Given our distinct phenome fingerprinting across the prioritized *C. neoformans* proteins, we mapped the virulence-associated phenome of the *C. neoformans* mutant strains relative to WT and generated a fingerprint of genetic contribution to key virulence factor elaboration (Fig. 2E). From this phenome fingerprint, we defined two categories with fungal therapeutic potential: antifungal (i.e., strain defect in all tested phenotypic assays) and anti-virulence (i.e., defect in virulence-associated defects, excluding fungal growth^35,36^). For the antifungal category, a well-characterized copper transporter and antifungal target, *bim1*Δ (serving as confirmation of our approach) and the uncharacterized, *CNAG_05997*Δ, were defined (Fig. 2F). For the antivirulence category, eight candidates were defined, including the well-characterized SOD1 with roles in reactive oxygen species neutralization^37,38^ (serving as confirmation of our approach), as well as proteins of interest, including CipC (CNAG_03007), an uncharacterized protein with a putative role as a biomarker of cryptococcal infection^18,39^ and virulence-associated roles in *A. fumigatus*^32,33,40,41^ (Fig. 2G).

### In vivo cryptococcosis model confirms antifungal and antivirulence proteins and reveals functional roles for CipC in cell wall stress resistance and immune modulation

Given the *in vitro* defects in classical virulence factor production and reduced LDH from macrophage upon infection, we assessed the classification of the candidates using an *in vivo* model of cryptococcosis for host survival and fungal burden^42–44^. For the *in vivo* models, the WT, two independent gene deletion strains, and a genetic complement strain were assessed. For *CNAG_05997*Δ, defined as a putative antifungal, we confirmed this designation with a significant increase in host survival (Fig. 3A) and significant reduction in fungal burden at primary (i.e., lungs) and secondary (i.e., brain, spleen) infection sites (Fig. 3B). These findings align with recent reports linking fungal pathogenesis to the mitochondria and electron transport chain^45,46^, highlighting the potential for CNAG_05997 as a novel putative druggable target. Next, we moved to the antivirulence category and given the observed role of CipC in fungal virulence but lacking a defined role in *C. neoformans*, we performed an *in vivo* cryptococcal infection assay. Here, we observed a significant increase in host survival (Fig. 3C); however, fungal burden was comparable to WT across the organs (Fig. 3D). These findings support the connection between energy production and antifungal potential, as well as suggest an important role for CipC in modulating the host response but without a role in fungal dissemination.

**Fig. 3:**
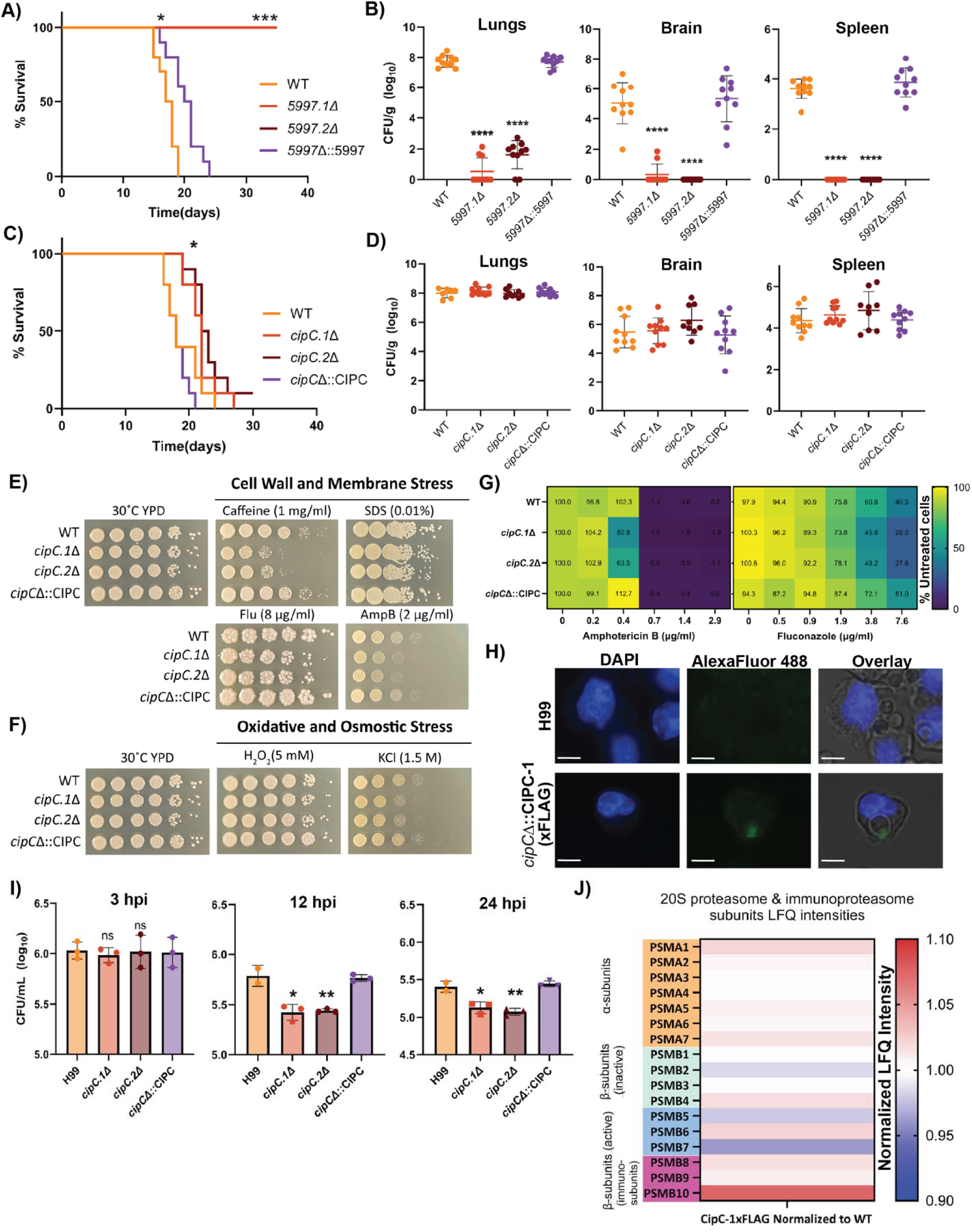
Evaluation and characterization of antifungal and antivirulence target. **A)** Survival curve for murine model of cryptococcosis performed with WT and two independent mutants of *CNAG_05997*Δ. **B)** Fungal dissemination profiles assessed by quantification of fungal burden from lung, brain, and spleen for *CNAG_05997*Δ. **C)** Survival curve for murine model of cryptococcosis performed with WT and two independent mutants of *cipCΔ*. **D)** Fungal dissemination profiles assessed by quantification of fungal burden from lung, brain, and spleen for *cipC*Δ. Experimental groups of mouse survival and fungal dissemination assays following inhalation model of cryptococcosis completed with n=10 female BALB/c mice. Differences in survival were statistically tested using a log-rank (Mantel-Cox) test (*, *P* ≤ 0.05; ***, *P* ≤ 0.0001). Differences in fungal burden statistically tested using Students *t-*test (*, *P* < 0.01; ****, *P* ≤ 0.0001). **E)** Cell wall and membrane stress susceptibility of *cipC*Δ growth on YPD supplemented with caffeine (1 mg/ml), SDS 0.01%, amphotericin B (2 µg/ml), and fluconazole (8 µg/ml), compared to YPD. **F)** Osmotic and oxidative stress susceptibility of *cipC*Δ growth on YPD supplemented with H_2_O_2_ (5 mM) and KCl (1.5 M) compared to YPD. Serial dilutions of strains were spotted on YPD supplemented with stressor and incubated at 30°C for 2-5 days. Experiment completed in biological duplicate and technical duplicate. **G)** Microdilution assay of amphotericin B sensitivity to strains at concentrations of 0-2.9 µg/mL and fluconazole sensitivity to strains at concentrations of 0-7.6 µg/mL. Cell density was measured at OD_600nm_ and represented as percentage of growth of the untreated control, corresponding percentage of growth labeled in each individual heat map cell. Experiment completed in biological quadruplicate and technical duplicate. **H)** CipC localization during *C. neoformans* infection of macrophages with *cipC*Δ::CIPC-1x FLAG strain detected during 3 h co-culture (MOI 100). Images captured with DAPI and Alexa Fluor 488 (FLAG). Scale bar = 5.1 µm. **I)** Engulfed fungal cells were quantified by co-culturing macrophages with *C. neoformans* strains for 3 h at an MOI of 10. Phagocytosed cells were enumerated for CFUs following PBS washes of infected macrophages. Intracellular fungal burden quantified following initial 3 h co-culture and PBS washes to remove extracellular and non-adhered cells, followed by maintenance in fresh DMEM. At 12 and 24 hpi, lysed macrophages were plated for CFUs. Statistical analysis using Students *t-*test (*, *P* ≤ 0.05; **, *P* ≤ 0.01). Experiment completed in biological triplicate. **J)** Normalized LFQ intensities (CipC-FLAG vs. WT) of host 20S proteasome and immunoproteasome subunits identified from CipC-1xFLAG interactome. Co-immunoprecipitation experiment completed in biological quadruplicate.

Given this gap in current knowledge, we pursued further functional characterization of CipC within the present study. Considering the conservation of CipC across fungal pathogens (Fig. S4A, S4B) and involvement in fungal stress responses (e.g., mycotoxin production, pathogenesis, growth conditions^32,33,41,47–49)^, we previously performed proteome profiling between WT and *cipC*Δ strains^50^. These data identified 25 fungal proteins with significant increases in abundance in the deletion strain, suggesting a connection with CipC production, including transcriptional regulators, Rho GTPase activator, and a peptidase, with diverse roles in virulence factor production (Table S3). To further assess this connection between CipC and fungal virulence and stress response, we performed plate sensitivity assays to caffeine and SDS, well as current antifungals that target fungal membranes, amphotericin B, and fluconazole and observed a reduction in growth for the *cipC*Δ strains (Fig. 3E). We also explored differences in oxidative and osmotic stress responses to KCl and H_2_O_2_ and observed a reduction in growth for the *cipC*Δ strains (Fig. 3F). Given the rising rates of antifungal resistance^51,52^ and the demonstrated increased susceptibility of *cipC*Δ to fluconazole and amphotericin B, we performed a limiting dilution assay. Here, we observed a significant increase in susceptibility to amphotericin B at 0.4 µg/mL and to fluconazole at 7.6 µg/mL for the *cipC*Δ strains compared to WT (Fig. 3G; Fig. S5). Together, these findings suggest new defined roles for CipC as a virulence determinant at the periphery of *C. neoformans* cells with dual roles in fungal protection and host immune modulation.

To further explore the putative role of CipC in host immune modulation, we performed immunofluorescence microscopy of macrophage infected with WT and CipC-Flag tagged cells. Here, we confirmed production of CipC during macrophage infection, with suggested localization to the fungal cell membrane and a potential interaction point within macrophage (Fig. 3H). Next, we assessed the intracellular susceptibility of *cipC*Δ within macrophages by quantification of fungal burden at 3, 12, and 24 hpi. We determined that deletion of *cipC* did not interfere with initial fungal uptake by macrophages; however, assessment of engulfed fungal cells at 12 and 24 hpi indicated significantly increased susceptibility to macrophage killing (Fig. 3I). These data further suggest a role for CipC in immune cell modulation.

To gain a better understanding into putative interactions between CipC and macrophages during infection, we performed a co-immunoprecipitation assay coupled with mass spectrometry. Production and detection of CipC was confirmed by immunoprecipitation (Fig. S6) and 21 host proteins were exclusively associated with the CipC-FLAG samples (Table S4). Notably, we observed an interaction between CipC and PSMB10, an immune-inducible subunit of the 20S proteasome induced upon infection involved in MHC class I antigen presentation^53^. To further explore this connection, we assessed potential interaction between CipC and other 20S proteasome and immunoproteasome (17 subunits^53^) and reported increased abundance for 14 of the subunits, further supporting a role for CipC in immune modulation (Fig. 3J). Together, these data showcase the role of CipC in *C. neoformans* toward resistance of cell wall stress, susceptibility to antifungal agents targeting the fungal membrane, localization to the membrane, along with immune system modulating interactions.

### cipCΔ-derived EVs feature defining sterol content and protein composition

To explore a connection across CipC-associated immune modulation, cell wall sensitivities, localization to the fungal cell membrane, and previous identification of CipC within cryptococcal extracellular vesicles (EVs) ^40^, we quantified differences in EV production between the strains. By quantification of ergosterol content of EVs derived from WT or *cipC*Δ strains, we observed significantly elevated (>2-fold) increase in ergosterol from *cipC*Δ derived EVs (Fig. 4A). Such a drastic difference suggests dysregulated biogenesis or exportation of EVs from the fungal cell upon *cipC* deletion. As EVs are engulfed and influence the infection dynamics of macrophages^54,55^, we evaluated if *cipC*Δ-derived EVs elicit an altered immune response. We assayed differences in macrophage cytotoxicity upon exposure to EVs derived from WT and *cipC*Δ strains and did not observe a significant difference compared to untreated macrophage (Fig. 4B). Next, we co-cultured macrophages with *C. neoformans* WT in the presence of WT-or *cipC*Δ-derived EVs and quantified intracellular burden levels. We determined that the presence of the EVs significantly decreased intracellular fungal burden at initial exposure (Fig. 4C), suggesting lowered phagocytosis of *C. neoformans* cells upon exposure to EVs. We assessed fungal burden again at 24 hpi (Fig. 4D) and confirmed a maintained significant reduction in fungal burden within macrophages exposed to EVs. Notably, the origin of the EVs (WT-or *cipC*Δ-derived) did not influence intracellular fungal burden levels. These data suggest that exposure of macrophage to EVs modulates the ability of macrophages to phagocytose *C. neoformans* cells.

**Fig. 4:**
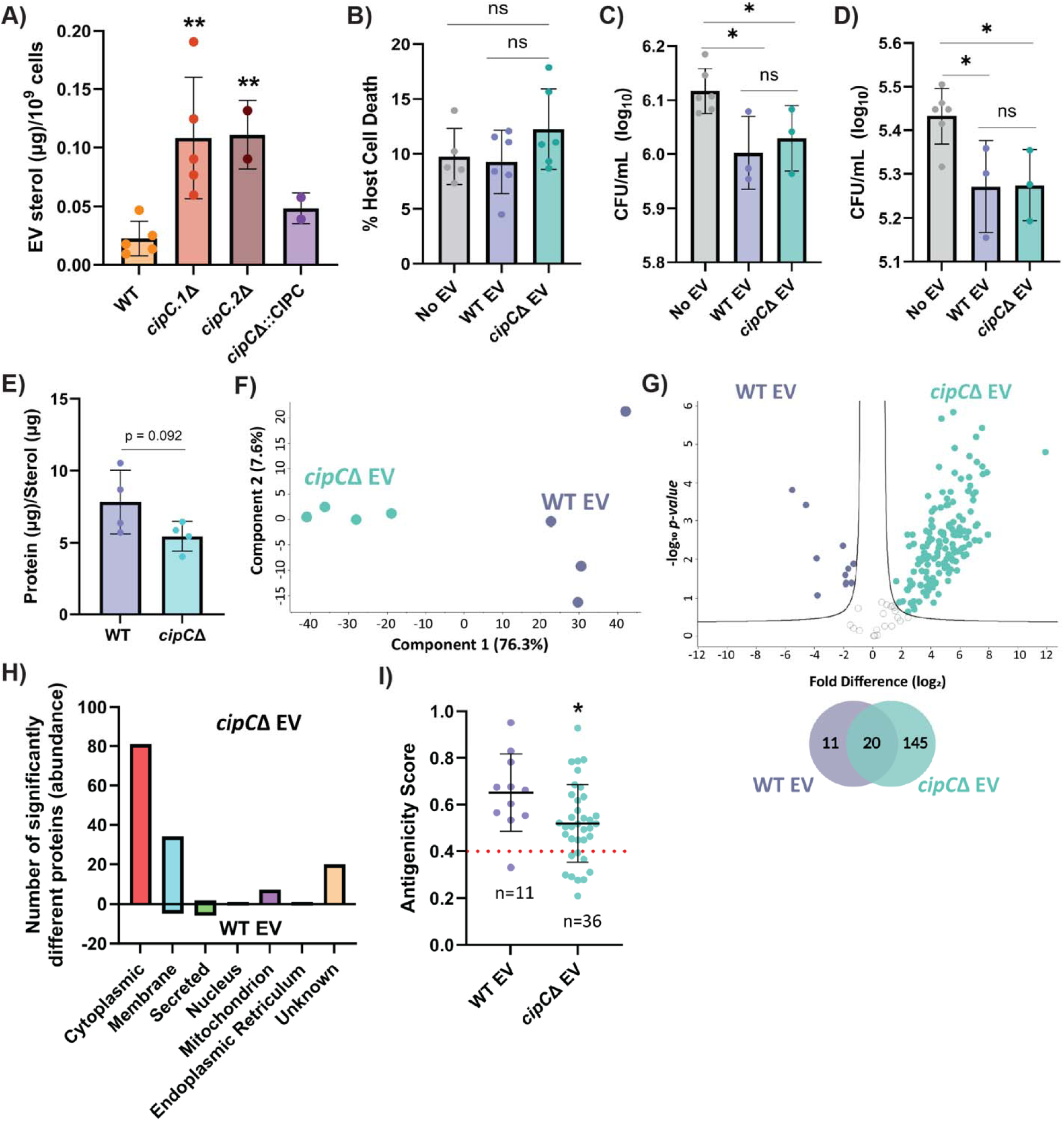
*cipC*Δ-derived EVs feature altered sterol and proteome profiles. **A)** EV production from *C. neoformans* strains on YPD solid agar at 30 °C; sterol quantity expressed per 10^9^ cells for each strain. Statistical analysis using Students *t-*test (**, *P* ≤ 0.01). **B)** Host cell cytotoxicity following treatment with 10 µg of WT and *cipC*Δ-derived EVs compared to PBS control. Treatment applied to BALB/c immortalized macrophages in serum-free DMEM for 3 h at 37 C and 5% CO_2_ followed by quantification of LDH release. **C)** During co-culture of *C. neoformans* WT with macrophages (MOI 10), 10 µg EVs (WT and *cipC*Δ-derived) were added at the same time as fungal cells in serum-free-DMEM. Phagocytosed fungal cells were quantified by CFUs at initial 3 h co-culture and PBS washes, followed by maintenance in fresh DMEM. **D)** At 24 h.p.i, lysed macrophages were plated for CFU quantification to assess affect of initial EV exposure on long-term infection dynamics. Statistical analysis using Students *t-*test (*, *P* ≤ 0.05). **E)** Total protein content of purified EVs normalized to sterol content, representative of amount of EV molecules per purification. Statistical significance tested using Student’s *t-*test, ns = not significant. Experiment completed in biological triplicate and technical duplicate. Values shown are representative of technical replicates from one independent assay. **F)** Principal component analysis for WT-and *cipC*Δ-derived EVs. **G)** Volcano plot comparing EV-associated proteins between WT and *cipC*Δ strains; Venn diagram depicts the number of significantly different proteins identified within each condition. Students t-test, *P* < 0.05; FDR = 0.05; S_0_ = 1. **H)** Distribution of GOCC terms between significantly different EV-associated proteins with significant increases or decreases between WT and *cipC*Δ origin strain shown. **I)** Antigenicity score of significantly different EV-associated proteins from corresponding origin strain classified with the GOCC term ‘membrane’ or ‘secreted’. Scores calculated using VaxiJen 2.0 Antigen Predictor. Statistical analysis of antigenicity scores completed using Students *t-*test (*, *P* ≤ 0.05). Proteomics experiment performed in biological quadruplicate.

As we observed differences in ergosterol production between WT-or *cipC*Δ-derived EVs but no difference in macrophage phagocytic response, we aimed to assess differences in EV protein cargo between the strains. Quantification of total protein content per EV sterol content (normalized) value showed a subtle decrease for *cipC*Δ-derived EVs but not to a significant level (Fig. 4E). We performed proteome profiling of EVs derived from WT or *cipC*Δ strains, and we observed clear separation between the EV source as component 1 at 76.3% and replicate reproducibility as component 2 at 7.6% (Fig. 4F). An assessment of significant changes in protein abundance clearly defined a modulated proteome within EVs derived from the *cipC*Δ strain with a significant increase in abundance of 145 proteins and a significant decrease in abundance of 11 proteins (Fig. 4G; Table S5). We categorized the significantly different proteins based on GOCC and reported an elevation of *cipC*Δ-derived EV proteins associated with cytoplasmic, membrane, mitochondrion, or unknown components (Fig. 4H). Given the distribution of protein functions within the EVs and our hypothesis that CipC modulates the host immune response, we applied the antigen prediction server, VaxiJen^56^, to predict the antigenicity of significantly different EV-associated proteins with roles in extracellular exposure (i.e., membrane, secreted) to the host. We observed a significantly higher antigenicity profile for WT-derived EV proteins, including virulence-associated cryptococcal proteins (e.g., cytokine inducing-glycoprotein^57^, chitinase^58^, immunoreactive mannoprotein MP88^59,60^) compared to *cipC*Δ-derived EVs (Fig. 4I). However, an enrichment in the total number of antigenic proteins was noted for *cipC*Δ-derived EVs (27 proteins) relative to WT (10 proteins) (Table S6). These differences support altered protein cargo between the EVs with a higher putative exposure of host cells to diverse extracellular antigens in the absence of CipC.

### Antigenic properties of cipCΔ-derived EVs potentiates a role in immunization against diverse fungal pathogens

Based on the EV proteomic profiling, we proposed that albeit WT-derived EVs may contain a stronger antigenicity score, the culmination of *cipC*Δ-derived EVs diverse and abundant antigen profile may produce a more robust, broad-spectrum protective immune response. To test this concept, we compared the immunoreactivity of WT-and *cipC*Δ-derived EVs using an established EV vaccination regime against cryptococcosis (Fig. 5A)^40^. With this triple-dose vaccination pilot, we immunized BALB/c mice with 10 µg EV-protein dosages via intraperitoneal injections, with the control group injected with only PBS. Following immunizations, we evaluated the anti-EV-antibody response in mouse sera with the appropriate loading control (Fig. S7). We determined that vaccination with *cipC*Δ-derived EVs resulted in a dramatic increase in antibody production against WT-EV molecules compared to WT-EV vaccination, whereas the PBS control appropriately displayed negligible immunoreactivity (Fig. 5B). Semi-quantification of the antibody response significantly corroborated the increased *cipC*Δ-derived EV immune response (Fig. 5C). These data further support the hypothesis that the increased combination of antigens present on *cipC*Δ-derived EVs elicit an advantageous robust immune response.

**Fig. 5:**
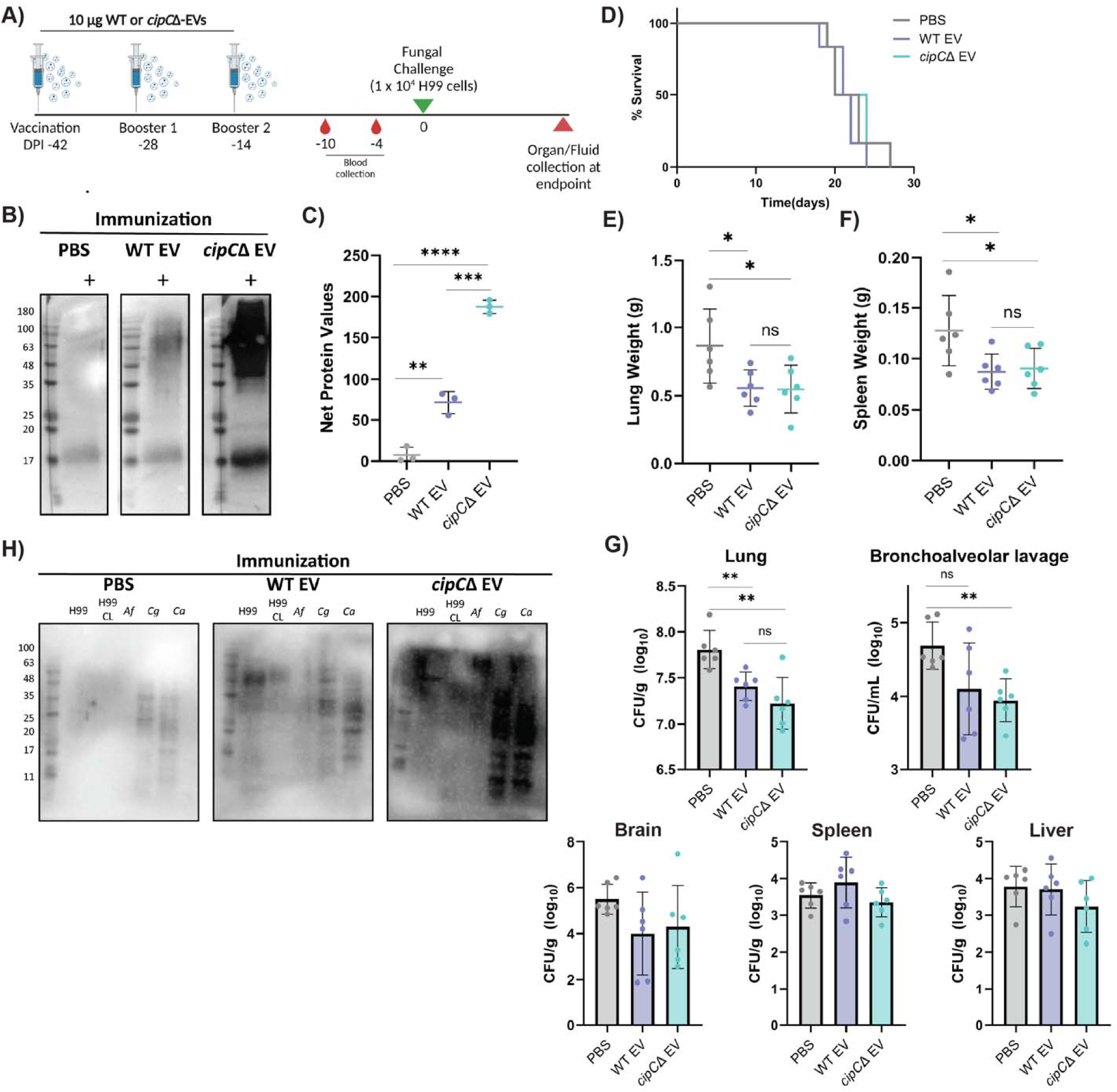
Immunization with *cipC*Δ-derived EVs leads to increased serological reaction. **A)** Experimental design of vaccination regimen and H99 challenge. Female BALB/c mice (*n* = 6 per group) were immunized by intraperitoneal injections of 10 µg EVs isolated from WT or *cipC*Δ strain in 100 µl of PBS. Control mice were injected with 100 µl PBS, followed by intranasal inoculation of *C. neoformans* WT. **B)** Western blot of WT-derived EVs probed by sera from nonimmunized and immunized mice. **C)** Quantification of serological response to EVs across vaccination regimes. Net protein values obtained from quantification of protein bands in Western blot analysis using ImageJ software. Experiment completed in technical triplicate. Statistical analysis using Students *t-*test (**, *P* ≤ 0.005; ***, *P* ≤ 0.0005; ****, *P* ≤ 0.0001). **D)** Survival rates of vaccinated animals with either WT- or *cipC*Δ-derived EVs and unvaccinated animals. **E)** Lung weights of vaccinated and unvaccinated mice infected with *C. neoformans* WT. **F)** Spleen weights of vaccinated and unvaccinated mice infected with *C. neoformans* WT. Statistical analysis using Students *t-*test (*, *P* ≤ 0.05). **G)** Fungal burden from lung, bronchoalveolar lavage, brain, spleen, and liver determined from homogenized tissues and quantification of CFUs. Statistical analysis using Students *t-*test (**, *P* ≤ 0.005). **H)** Cross-reactivity of serological reaction of immunized mice to other major human fungal pathogens, including, including a *C. neoformans* clinical isolate of strain H99 serotype A, *Aspergillus fumigatus* ATCC 204305, *Candida glabrata* BG87, and *C. albicans* SC3514. Experiment completed in technical duplicate.

To determine if the EV vaccines provide protection within a cryptococcosis murine model, we intranasally challenged mice with *C. neoformans* WT strain. With the tested immunization model, the EV-immunized mice displayed the same survival rates as nonimmunized mice (Fig. 5D); however, evaluation of fungal dissemination profiles and fungal burden levels showed significant reduction in lung and spleen weights from immunized mice; indicative biomarkers of lowered pulmonary cryptococcosis and inflammation^61,62^ (Fig. 5E, 55F). We observed significantly reduced fungal pulmonary burden associated with EV immunization, with the *cipC*Δ EV immunized mice lungs indicating a modestly lower burden than the WT-EV vaccinated mice (Fig. 5G). However, fungal burden from dissemination to secondary sites (i.e., brain, spleen, liver) were not significantly different upon immunization, although lower trends corresponding to immunization status were observed in the brain.

Finally, we explored if EV immunizations could mount a protective response against other common myotic infections as a potential pan-fungal vaccine candidate. Herein, a comparison of the anti-EV-antibody response of nonimmunized and immunized sera revealed substantially increased immunoreactivity from *cipC*Δ EV-immunized sera against tested fungal pathogens, including *C. neoformans* H99, *C. neoformans* clinical isolate, *Aspergillus fumigatus*, *Candida glabrata*, and *Candida albicans* (Fig. 5H). These findings support the potential of *cipC*Δ-derived EVs as a putative immunization source given the occurrence of common antigens expressed across multiple fungal pathogens that currently lack approved immunization strategies.

## Discussion

In this study, we developed an innovative pipeline driven by quantitative proteomics combined with phenome fingerprinting to identify and characterize novel potential fungal therapeutic targets. We focused on a critically important interface within cryptococcal disease: the interaction with macrophages. Despite the impressive previous characterizations of *Cryptococcus* and macrophages, few studies have reported a dual perspective of the interacting species’ proteomes. Our approach enabled robust identification and measurement of the host response to fungal infection, revealing immune suppression themes typical of *C. neoformans* virulence to diminish host defense and prolong fungal maintenance within the host^63^. Moreover, we defined a subset of 53 fungal proteins to be highly abundant when *C. neoformans* is in the presence of the host. Our strategy to prioritize proteins that are secreted or membrane-bound, uncharacterized, and associated with cryptococcal virulence prioritized a library of 15 candidate proteins for further investigation. Through a comprehensive assessment of protein functional roles in cell wall stress, immune modulation, and antigenicity, we revealed previously unreported immunization power of CipC in *C. neoformans* and further defined a putative new role in host immunization against diverse human fungal pathogens.

This study offers some rationale limitations that may influence complete elucidation of the *C. neoformans* infectome. For example, our decision to dissect the interaction of *C. neoformans* with monocytic-derived rather than tissue-resident macrophages arises from the fungus’ mobility and plasticity to exploit circulating macrophages for brain and CNS dissemination; thus, fungal candidates of interest are more likely to be employed for virulence across all cryptococcal infection stages^64–66^. Furthermore, our experimental infectome setup (i.e., MOI inoculum, co-culture time) is a narrow perspective of cryptococcal infection and does not fully encompass this exhaustive interaction. However, in comparison to recent cryptococcal-host proteome characterizations featuring diverse infection conditions (i.e., opsonization, bacterial co-infection, murine models), findings indicate a common core infectome. For instance, CipC, was identified as a putative biomarker of cryptococcal infection from the blood^39^ and spleen with mapped unique protein production patterns ^42^ and as a chronic infection-associated candidate protein^63^. These observed frequent fungal profile signatures for CipC highlight the dependability and robustness of our pipeline.

Many *C. neoformans* virulence traits operate distinctly at the fungal cell surface and extracellular environment (e.g., capsule, melanin, and enzyme production); thus, our consideration for extracellular exposed fungal proteins in our selection criteria supports the potential to define fungal-host interactions and virulence elaboration^8,67^. Our systematic fingerprinting of the candidate fungal strains revealed impressive divergence of virulence phenomes compared to WT, leading to benchmarking of two therapeutic categories: antifungal and anti-virulence. Assessment of selected therapeutic candidates in murine models of cryptococcosis validate our approach to identify antifungal targets and exposed the importance of antivirulence targets in immune system regulation. The antifungal candidate, CNAG_05997, holds remarkable promise as a potential novel antifungal target based on the disruption of all major fungal virulence factors and nearly complete eradication of the fungus in the murine infection model. Moreover, CNAG_05997 features predicted localization to the mitochondria, proposing essential survival functions. Many antifungal drugs with approved clinical use exclusively target fungal cell walls and plasma membrane components, and the current development of new agents focuses on the derivatization of existing compounds^6^. Therefore, expanding the repertoire of antifungal targets is critical to reduce redundant mechanisms of action^68^. A handful of mitochondria-targeting lead compounds in development hold promise for inhibiting this essential organelle and its pivotal roles in stress response, morphogenesis, virulence, and drug resistance^69–72^. Considering CNAG_05997 features minimal overlapping mammalian homology, our data supports this protein as a fungal-specific mitochondrion protein and is a desirable antifungal target; however, further investigations are required to delve into the protein’s druggability.

Our prioritization of CipC as the infection-relevant representative candidate from our phenome fingerprinting derives from the undiscovered promise of this target. Interestingly, CipC-like homologs are exclusively found in fungi and are more commonly reported in filamentous fungi rather than yeast and its presence in *A. fumigatus* hyphal-morphotype, supports a role for aspergillosis invasive growth ^32^. One of the major differences between yeast and hyphae forms is cell wall scaffolding^73^. Thus, we provide multiple lines of evidence supporting the importance of CipC for cell wall and plasma membrane integrity. Moreover, as cryptococcal EVs and immunogenic proteins have promise as immunization candidates, we reasoned that the adjustment of proteins on *cipC*Δ-derived EVs could potentially increase protection in a vaccine model^40,74–76^. We found that *cipC*Δ-EVs showcased modestly higher cytotoxicity to macrophages, indicating potential macrophage stimulation, and both EV types resulted in significantly less intracellular fungal burden. Whether this results from enhanced macrophage killing or activated fungal evasion techniques remains to be determined. Regardless of the EV origin strain, the immunization model tested was insufficient to prolong murine survival. This lack of protection may be attributed to the hypervirulence of the H99 strain, requiring additional dosage optimizations compared to the reference trial conducted with KN99α^40,77,78^; however, notably, the dramatic increase of immunoreactivity from the sera of *cipC*Δ-EV immunized mice without the use of adjuvants or carriers highlights a more robust antigen-specific immune response than WT-EV vaccination strategy.

Finally, given that a single conserved antigen can confer cross-protection to different fungal pathogens, as illustrated by the NXT-2 and calnexin vaccine candidates^79,80^, we proposed that the dramatic increase of cargo proteins packaged into *cipC*Δ-EVs, share amplified conservation with other fungal pathogens. For instance, a vaccine-based immunopeptidomic approach to identify *Histoplasma capsulatum* vaccine epitope candidates revealed promising epitope conservation with eight medically important fungi, including intracellular proteins Hsp60, ATP synthase subunit alpha, and elongation factor 1-gamma, all of which we identified to be significantly abundant within *cipC*Δ-EVs^81^. We observed an enhanced antibody signal from the sera of *cipC*Δ EV-immunized mice, capable of binding to epitopes from whole cell lysates of four major fungal pathogens. It is possible that the epitopes recognized by the protective antibodies elicited by *cipC*Δ EV-immunizations are recognizing various conserved epitopes across both the filamentous and yeast pathogens assessed. The translational significance of these findings is critical considering vaccine requirements to contain sufficient antigens to elicit a broad immune response; however, an abundance of antigens can correlate with heightened side effects when vaccinating an immunocompromised population, requiring careful consideration.

## Conclusion

In summary, this study introduces a scalable, unbiased infectome–phenome discovery framework that bridges high-resolution proteomics with functional phenotyping to systematically uncover fungal drivers of infection. By resolving coordinated host immune suppression and fungal virulence programs during *Cryptococcus neoformans*–macrophage interactions, this platform enables principled prioritization of infection-associated proteins with therapeutic relevance. Functional stratification of candidate targets into antifungal and antivirulence categories, coupled with *in vivo* validation, underscores the power of this approach to move beyond descriptive omics toward actionable biological insight. The identification of CipC as a conserved virulence-associated factor further reveals a previously underappreciated role for fungal EVs in shaping host immune responses and antigenic landscapes. Collectively, these findings provide a versatile framework for interrogating fungal pathogenesis and accelerating the discovery of effective intervention strategies applicable across a broad spectrum of invasive fungal diseases.

## Materials and Methods

### Fungal strains and growth conditions

*Cryptococcus neoformans* var. *grubii* wild-type (WT) strain H99 (serotype A) was used for all analyses and as a background strain for mutant construction. *C. neoformans* WT strain was cultured on yeast extract-peptone-dextrose (YPD) agar plates (2% dextrose, 2% peptone, 1% yeast extract, 1.5% agar), mutant strains and complement strains were cultured on YPD supplemented with 100 μg/mL nourseothricin (NAT) and 200 μg/mL hygromycin B (HYG), respectively, at 30 °C, unless otherwise stated. For routine growth, all fungal strains were cultured in YPD media at 30 °C with shaking at 200 rpm, unless otherwise stated. For *in vitro* growth cultures, WT was grown in YPD media to mid-log phase (12 h) at 37 °C and collected for proteome and secretome analysis.

### Bone marrow-derived macrophage collection and infection

For the infectome analysis, BALB/c WT immortalized bone marrow-derived macrophages (BMDM) (generously prepared and provided by Dr. Felix Meissner, Max Planck Institute of Biochemistry, Germany) were prepared for cryptococcal infection as described previously^82^. Macrophages were maintained in Dulbecco’s modified eagle’s medium (DMEM) supplemented with 10% (v/v) heat-inactivated fetal bovine serum (FBS), 2 mM Glutamax, 1% sodium pyruvate, 1% L-glutamine, and 5% pen/strep. BMDMs were seeded in 6-well plates (i.e., 1 x 10^6^ cells/well) in BMDM- DMEM media without pen/strep overnight at 37 °C, 5% CO_2_. Fungal cells were grown in YPD media to mid-log phase (same cultures used for *in vitro* growth cultures as described above), centrifuged and the supernatant collected for secretome processing. The cell pellet was washed twice in phosphate-buffered saline (PBS), an aliquot for fungal *in vitro* proteome processing was collected, and fungal cells were diluted in BMDM-DMEM media.

BMDMs were infected at a multiplicity of infection (MOI) of 100:1 (fungal:macrophage) for 3 h at 37 °C, 5% CO_2_. Control samples consisting of uninfected macrophages were maintained in BMDM-DMEM without pen/strep to correspond with the progression of fungal infection. Infected and uninfected BMDM cells were washed twice with PBS to remove non-phagocytosed fungal cells and collected for proteome analysis. The BMDM-fungal infection assay was performed using biological triplicates.

### Proteomic sample preparation

Proteomic profiling, including proteome, secretome, and infectome, was performed as previously described with minor modifications^63,83^. Briefly, PBS-washed cell pellets (fungal and macrophage) were resuspended in 100 mM Tris-HCl (pH 8.5) with a protease inhibitor cocktail tablet (PIC; Sigma-Aldrich), followed by probe sonication (Thermo Fisher Scientific) in an ice bath. Sodium dodecyl sulfate (SDS) (2% final) was added, followed by reduction with 10 mM dithiothreitol (DTT) at 95 °C for 10 min with 800 rpm. Next, alkylation was performed with 55 mM iodoacetamide (IAA) at room temperature in the dark, and acetone precipitation (80% final) overnight at-20 °C. Samples were collected and washed twice in 80% acetone, dried, and resuspended in 8 M urea-40 mM HEPES for protein quantification using bovine serum albumin (BSA) tryptophan assay^84^. Samples were diluted to 2 M urea with 50 mM ammonium bicarbonate (ABC), normalized to 100 µg protein, and digested with LysC-trypsin (Promega [protein/enzyme ratio, 50:1]). For secretome samples, the fungal-YPD supernatant was filtered (0.22 µm syringe filter tip), reduced with 10 mM DTT (30 min, room temperature), alkylated with 55 mM IAA (20 min, in the dark), and digested with LysC-trypsin overnight as described above. Digestion was stopped with 10% (v/v) trifluoracetic acid (TFA), and acidified peptides were purified using Stop-And-Go extraction tips (Stage tips; containing three layers of C_18_) to desalt and purify following standard instructions^85^.

### LC-MS/MS

Peptides were initially resuspended in Buffer A (2% acetonitrile [ACN], 0.1% TFA) for liquid chromatography-tandem mass spectrometry (LC-MS/MS) analysis. Peptides were subjected to nanoflow liquid chromatography on an EASY-nLC system (Thermo Fisher Scientific, Bremen, Germany) on-line coupled to an Q Exactive HF or QExactive 240 quadrupole orbitrap mass spectrometer (Thermo Fisher Scientific). Samples were loaded onto a 50 cm column with 75 µmm inner diameter, in-house packed with 3 µm reversed-phase silica beads (ReproSil-Pur C_18_-AQ, Dr. Maisch GmbH, Germany). Separated peptides were electrosprayed into mass spectrometer with a linear gradient of 5% to 50% ACN in 0.5% acetic acid over 120 min at a constant flow of 300 nL/min, followed by a wash with up to 95% ACN. The mass spectrometer operated in a data-dependent acquisition mode alternating between one full scan and MS/MS scans of abundant peaks (i.e., Top 15 method). Full scans (*m/z* 300-2000) were acquired in the Orbitrap analyzer with a resolution of 60,000 at 100 *m/z* or 120,000 at 200 *m/z*.

### Data Processing and MS bioinformatics

Mass spectrometry raw data files were processed using MaxQuant software (v. 1.6.2.10)^86^. The derived peak list was incorporated with the Andromeda search engine against the reference *C. neoformans* var. *grubii* serotype A (strain H99/ATCC 208821) proteome (7430 sequences; Oct. 2018) and *Mus musculus* (55,462 sequences; Oct 2018) from Uniprot^87^. The parameters were as follows: trypsin enzyme specificity allowing for a maximum of two missed cleavages, seven amino acid minimum required peptide length, fixed modification of cysteine carbamidomethylation, variable modifications of methionine oxidation and N-acetylation for proteins. A target-decoy approach filtered peptide spectral matches with a false-discovery rate (FDR) of 1%, with a minimum of two peptides required for protein identification. Relative label-free quantification (LFQ) and match between runs enabled, and the MaxLFQ algorithm used a minimum ratio count of 1^88^. Split by taxonomic ID enabled when two proteome FASTA files were inputted in the same experiment (e.g., *C. neoformans* and *M. musculus*).

Statistical analysis and data visualization of MaxQuant-processed data was performed using Perseus software (v. 1.6.2.2)^89^ and ProteoPlotter^90^. First, data were prepared by filtering for reverse hits to the database, contaminants, and proteins only identified by site, followed by log_2_ transformation of LFQ intensities. Filtering for valid values (valid-value filter of 3 in at least one group) and missing values were imputed from the normal distribution (width, 0.3 standard deviations (SD); downshift, 1.8 SDs). Significant differential abundance of proteins identified by Student’s *t*-test (*P* ≤ 0.05) with multiple-hypothesis testing correction using the Benjamini-Hochberg FDR cutoff at 0.05 with S_0_ = 1^91^. STRING database was used for protein-protein interaction network mapping as described at https://string-db.org/. Protein-protein interaction networks generated with STRING basic settings and medium confidence (i.e., 0.4). STRING enrichment analysis completed in the statistical background of the whole genome of *M. musculus*. The 1D annotation enrichment was performed as previously described^92^.

### Gene deletion and complementation

Generation of all mutant and complementation strains were obtained by directly introducing PCR product into H99 or mutant strain, respectively, by biolistic transformation^93^. For gene deletion constructs, linear constructs (1-1.5kbp) containing NAT resistance cassette (generously provided by Dr. J.P. Xu, McMaster University) linked to homologous 5’ and 3’ flanking regions of coding gene from H99 strain were generated by double-joint PCR as previously described^29^. Stable transformants were selected on YPD-NAT (100 µg/mL) agar, confirmed by diagnostic PCR, and two independent mutants per deletion of coding gene were selected for phenotypic characterization. Mutant strains selected for therapeutic candidate functional characterization (i.e., murine survival assay) were confirmed for single insertion of PCR product into the genome by Southern blot analysis probing for NAT cassette.

For complementation strains, the constructs were generated by amplifying the coding genes endogenous promoter (∼1 kbp upstream), open reading frame with a 1x-FLAG added to the 3’ end of coding gene, terminator (∼500 bp downstream), HYG resistance cassette, and 3’ flanking region (∼1 kbp) followed by Gibson assembly into pSDMA58 (generously provided by Dr. James Fraser, University of Queensland (Addgene plasmid # 67942; http://n2t.net/addgene:67942; RRID:Addgene_67942)). The constructs were amplified from vector to obtain transformation PCR products. Stable transformants were selected on YPD-HYG (200 µg/mL) agar, confirmed by diagnostic PCR, western blot, and Southern blot analysis probing for HYG cassette. Primers for genetic deletion and restoration (Table S7).

### Antifungal, antivirulence, infection-relevant candidate selection

Antifungal criteria filters for candidate deletion strains with phenotypic defects featuring normalization values <1.0 across all phenotypic assays (i.e., capsule size, capsule:cell ratio, melanin pigmentation, growth at 30 and 37 °C, and LDH release). Antivirulence criteria filters for candidate deletion strains, excluding candidates filtered in the antifungal category, for any virulence-related defect (i.e., capsule, melanin, thermotolerance, LDH release) with a normalization value <0.85 across both independent positive mutant strains were prioritized^94^.

### Phenotypic virulence-associated assays

To analyze fungal growth and thermotolerance of *C. neoformans* strains, fungal cells were inoculated in YPD media and grown overnight at 30 °C. The OD_600nm_ of overnight cultures was diluted to a final concentration 0.02 in yeast-nitrogen-base (YNB) medium with amino acids (BD Difco, Franklin Lakes, NJ) supplemented with 0.05% dextrose and incubated at either 30 °C or 37 °C. OD_600nm_ measurements were obtained on BioTek HM1 plate reader every 15 min over 72 h. To supplement liquid thermotolerance assays, dilution plate assays were performed as previously described^29^. Briefly, cells were collected, counted, and resuspended in YNB followed by a tenfold serial dilution (10^6^ cells/5 µl) plated on YNB agar plates. Plates were incubated at either 30 °C or 37 °C for 3 d, with images taken every 24 h.

Polysaccharide capsule production was visualized for *C. neoformans* strains following growth overnight in YPD, followed by a 1:100 subculture in YNB overnight, cells were washed twice in low-iron media (LIM; prepared as previously described^95^), and inoculated into LIM 1:100 for 24 h at 37 °C. Capsule production was visualized using India ink dye (Hardy Diagnostics) on a Zeiss Axiovert microscope 200M equipped with a Hamamatsu C10600 Orca R^2^ camera at 100X magnification under differential interference contrast (DIC). Cell diameter and capsule thickness were measured from a minimum of 50 cells per strain using ImageJ software (https://imagej.nih.gov/ij/index.html). Relative capsule sizes are reported as the ratio of capsule thickness to cell diameter.

Melanin pigmentation was assessed for *C. neoformans* strains following growth overnight in YPD, followed by a 1:100 subculture in YNB overnight, and a final 1:100 subculture in minimal media (MM; Chelex® 100-treated (Bio-Rad) dH_2_O containing 29.4 mM KH_2_PO_4_, 10 mM MgSO_4_–7 H_2_O, 13 mM glycine, 3 μM thiamine, 0.27% dextrose) overnight at 30 °C. Cells were collected, counted, and resuspended in minimal media (MM) followed by spot plating (10^5^ cells/5 µl) on MM agar plates supplemented with 1 mM L-DOPA (Sigma-Aldrich). Spots of fungal cells were evenly spaced on agar to avoid cross-contaminated cell density melanization. Plates were incubated at either 30 °C or 37 °C for 3 d, with images taken every 24 h. Percentage of melanin pigmentation was quantified as previously described^96^. Briefly, each image was converted to 8-bit grayscale, and the spot gray value (0-255) was calculated from a set area (50 x 50 pixels) on ImageJ software. Percentage of pigment was calculated by: [(255 subtracted by the mean grayscale value for each spot)/255] x 100.

### Extracellular vesicle purification

EV isolation was performed as previously described with minor modifications^40,97^. One colony of each strain was inoculated into 5 ml of YPD media and cultivated for 24 h at 30 °C. Cells were collected, washed twice in 10 ml of sterile water, and diluted to 3.5 10^7^ cells/ml in water. Cell suspension was aliquoted (300 µl) and spread onto YPD plates containing 34 µg/mL chloramphenicol and incubated for 24 h at 30 °C. Six petri dishes were used for each EV isolation. Cells were gently removed with a cell scraper (VWR), resuspended in 30 ml of 0.22 µm-filter sterile PBS and pelleted at 5,000 g for 15 min at 4 °C. The supernatant was centrifuged again at 15,000 g for 20 min at 4 °C followed by filtration through a 0.45 µm syringe filter. The filtered supernatant was ultracentrifuged at 100,00 g for 1 h at 4 °C (Type 70 Ti fixed-angle rotor, Beckman Coulter). The resulting pellet was resuspended in 0.22 µm-filter sterile PBS and immediately stored at-80 °C. The total sterol quantity was measured by the Amplex™ Red Cholesterol Assay Kit (ThermoFisher, A12216) and total protein content measured by Pierce™ BCA Protein Assay Kit (ThermoFisher, 23225). WT and *cipC*Δ-derived EVs were prepared and subjected to LC-MS/MS analysis as outlined above.

### Macrophage infection assays

For virulence-associated macrophage infection assays, BALB/c WT immortalized macrophages were used as described above. Macrophages were seeded in 24-well plates at 0.05 10^6^ cells/well and grown until 70-80% confluence (approximately 48 h). *C. neoformans* strains were grown to mid-log phase in YPD at 30 °C, collected, and washed twice in PBS. Fungal cells were resuspended in DMEM without pen/strep and opsonized with mAb18B7 (1 µg:10^6^ cells) for 1 h at 37 °C and 5% CO_2_. Macrophages were infected at an MOI of 10 for 90 min at 37 °C, 5% CO_2_, followed by two PBS washes, and addition of fresh DMEM without pen/strep for 27 h. Macrophage cell death was quantified at 27 h post-inoculation (hpi) as described previously^98^. The percentage of cell death was determined by lactate dehydrogenase (LDH) quantity in culture supernatants quantified by CytoTox96® Non-Radioactive Cytotoxicity Assay (Promega) according to manufacturer’s instructions. LDH quantification was performed in biological triplicate, and the experiment was performed in technical duplicate or triplicate.

The infectivity of *cipC*Δ strains was assessed by macrophage uptake and burden during prolonged infection by quantification of intracellular colony-forming units (CFU). BALB/c macrophages were infected in 12-well plates with *C. neoformans* WT, *cipC*Δ and complement strains at an MOI of 10 for 3 h. Following co-culture, wells were washed with PBS to remove extracellular and non-adhered fungal cells, and samples were collected for an initial 3 h (i.e., phagocytic uptake) time point by lysing macrophages with 0.5% triton at room temperature for 10 min. Fresh DMEM was added to the remainder of the wells for sample collection at 12 and 24h. Collected lysed macrophages were serially diluted and plated on YPD, followed by incubation at 30°C for 48 h for CFUs. Experiment completed in biological triplicate.

The ability of the purified WT and *cipC*Δ-derived EVs to impact *C. neoformans* WT infection of macrophages was assessed by enumeration of CFUs and LDH, as described above.

Briefly, BALB/c macrophages were infected in 12-well plates with *C. neoformans* WT at an of MOI 10 in the absence or in the presence of 10 µg (protein content) of WT or *cipC*Δ-derived EVs in serum-free DMEM (SF-DMEM). Following 3 h co-culture, wells were washed with PBS and fresh DMEM added for 24 h of infection. After 3 h and 24 h of infection, CFU and LDH measurements were collected. Experiment completed in biological triplicate and technical duplicate.

### Stress dilution plate assay

The sensitivity of *cipC*Δ to various stressors was analyzed by stress dilution plate assays performed as previously described^99,100^. All stressors were supplemented to YPD agar plates as follows: osmotic stressor, KCl (1.5 M); cell wall stressors, caffeine (1 mg/mL), SDS (0.01%), amphotericin B (2 µg/mL), fluconazole (8 µg/mL); and oxidative stressor, H_2_O_2_ (3 mM). *C. neoformans* strains were grown to mid-log phase in YPD at 30 °C, serial diluted ten-fold (10^6^ cells/5 µl) on supplemented YPD plates and incubated at 30 °C. Images were taken every 24 h for 5 d. Experiment completed with two biological replicates and technical replicates.

### Microdilution antifungal sensitivity assay

Determination of the MICs needed to inhibit *cipC*Δ growth relative to that of vehicle-treated cells were performed as previously described with minor modifications^101^. Briefly, cells were grown overnight in YPD at 30 °C and diluted to an OD_600nm_ of 0.01 in YNB. Treated and control cells were statically incubated in clear, round-bottom 96-well plates at 30 °C for 48 h, and the OD_600nm_ measured. Sensitivity to antifungal drugs were determined using concentrations over a five-dilution series of 0.18 to 2.85 µg/mL for amphotericin B and 0.48 to 7.6 µg/mL for fluconazole. Each assay done in biological quadruplicate and technical duplicate. Reported values are representative of independent assays. Data reported as the percentage of cell density relative to that of untreated cells per respective strain.

### Immunofluorescence

BALB/c WT immortalized macrophages were grown in FluoroBrite DMEM media (Thermo Fisher Scientific) supplemented with 10% FBS and 1% L-glutamine for 24 h prior to infection and throughout the infection protocol to minimize autofluorescence. Macrophages were infected at an MOI 100 with *C. neoformans* WT and *cipC*Δ::CIPC-1xFLAG strains for 3 h, following co-culture samples were washed and immediately fixed in 4% paraformaldehyde. Protocol for immunostaining was adapted from previously described methods^102,103^. Briefly, fixed cells were plated on 0.1% poly-L-lysine (Sigma-Aldrich) coated slides and incubated with lysing buffer (0.1 M sodium citrate, 1.1 M sorbitol pH 5.5) containing 10 mg/mL of lytic enzymes (Sigma, L1412) and a PIC and incubated at 30 °C for 2 h. Slides were submerged in 99% methanol, followed by 100% acetone, and then blocked with 2% goat serum, 2% BSA, and 0.1% saponin in PBS for 1 h. Blocked cells were incubated with 1:100 dilution of anti-FLAG M2 antibody (Sigma-Aldrich) for 1 h followed by 1 h incubation with 1:200 dilution of anti-mouse Alexa Fluor 488 (AF488; Invitrogen) and DAPI (10 µg/mL). Slides were washed between all incubation steps three times with PBS containing 0.01% Tween-20. Coverslips were mounted with a drop of *SlowFade^TM^* Gold antifade (Life Technologies). Slides were imaged using a Leica CTR5500B equipped with X-Cite series 120Q (Lumen Dynamics) and a pco.panda 4.2 M (Excelitas Technologies) camera through Volocity software ver. 6.6 (Quorum Technologies). A fixed exposure of 2.5 s was used to detect Alexa Fluor 488 bound to fungal cells, 9 ms for DAPI, and 59 ms for phase contrast. Experiment completed in biological triplicate. Production of CipC-FLAG measured across 86 fields of view with 85% of FLAG-labelled cells emitting fluorescence normalized against unlabelled WT cells.

### Co-immunoprecipitation

WT (negative control) and *cipC*::CIPC-FLAG strains infected BALB/c macrophages at an MOI 100 for 3 h, as described above. Following co-culture, wells were washed two times in PBS, and lysed in cold lysis buffer (150 mM NaCl, 50 mM Tris pH 7.5, 5% glycerol, 0.1% NP-40, PIC tablet), collected, and allowed to lyse further on ice for 10 min. Lysates were centrifuged at 4000 g for 5 min at 4 °C to pellet insoluble material. The supernatant was incubated with anti-FLAG Magnetic Agarose beads (Pierce) for 80 min with end-over-end rotation at 4 °C. Beads were washed twice with chilled wash buffer I (150 mM NaCl, 50 mM Tris pH 7.5, 0.25% NP-40) and four times with room temperature wash buffer II (150 mM NaCl, 50 mM Tris pH 7.5), followed by on-bead digestion as previously described^104^. Digested samples were purified by STAGE-tip and subjected to LC-MS/MS protocol outlined above.

### Murine model of cryptococcosis

Murine infection assays were performed according to the legislation and regulations of the Animals for Research Act and the Canadian Council on Animal Care. The animal models and procedures used have been approved under the Animal Utilization Protocol (AUP) 4193 at the University of Guelph.

(i) *Virulence*

For candidate functional characterization (i.e., CNAG_03007 and CNAG_05997) within *in vivo* assays, *C. neoformans* strains (i.e., WT, mutant, complement) were grown overnight in YPD at 30 °C and subcultured in fresh YPD to mid-log phase, collected, washed twice in PBS, and resuspended at 4 10^6^ cells/mL in PBS. Ten female BALB/c elite mice aged 7-9 weeks old (Charles River Laboratories, ON, Canada) were intranasally inoculated with 2 10^5^ cells (i.e., 50 µl of *C. neoformans* cell suspension) under isoflurane anesthesia. The mice were monitored daily for signs of morbidity and euthanized by isoflurane and CO_2_ inhalation at animal endpoint. Endpoint criteria include animal deterioration to specifications, including loss of 20% body weight, respiratory issues, lethargy, ruffled coat and hunched posture, or visible signs of neurological deficits. Organ and fluid collection of lungs, bronchoalveolar lavage, brain, spleen, and liver were collected upon termination. The collected tissues were weighed and homogenized in 1 ml PBS using a Bullet Blender Storm (Next Advance, Troy, NY, USA), serially diluted and plated on YPD agar supplemented with 32 µg/mL chloramphenicol (Sigma) and incubated for 48 h at 30 °C and assessed for CFUs. Tissues and fluids associated with *CNAG_05997*Δ infection were incubated for up to five days at 30 °C to account for the severe growth defect of the mutant strain and to allow adequate time for CFU observation.

(ii) *Immunization*

The EV-cryptococcal vaccination regime was performed as previously described^40^, with minor modifications. EVs were prepared for vaccination as described above. Six female BALB/c elite mice aged 7-9 weeks old (Charles River Laboratories, ON, Canada) were used for each group within the immunization experiment. The BCA method quantified protein concentration of the EVs for vaccination. Mice were vaccinated with EVs derived from WT, *cipC*Δ, or a non-EV control (i.e., PBS, non-immunized) three times, at 14 day-intervals with intraperitoneal injections (fixed protein amount of 10 µg in 100 µl PBS). Blood was collected from the submandibular veins of the mice 4 days and 10 d after the last immunization and tested for EV-specific antibody response by western blot^40,105^. All immunized and non-immunized control mice were intranasally infected with 1 10^4^ *C. neoformans* WT cells in 50 µl PBS. Animal morbidity and endpoint criteria were monitored as stated above. Organ and fluid collection of lungs, bronchoalveolar lavage, brain, spleen, and liver were retrieved and assessed for fungal burden as described previously.

### Western blot

(i) *Vaccine EV-antibody response*

Immunoblotting to test for antibody response of the immunized mice was performed as previously described^40^. Briefly, 10 µg protein aliquots of EVs (WT and *cipC*Δ-derived) were separated by 12% SDS-PAGE and transferred to a polyvinylidene fluoride (PVDF) membrane using a transfer apparatus according to manufacturer’s instructions (Bio-Rad). Membranes were blocked overnight in 5% BSA in Tris-buffered saline (TBS) (50 mM Tris, 150 mM NaCl [pH 7.5]) at 4 °C, washed two times with TBST (TBS, 0.05% Tween 20) and incubated with mouse sera at dilution 1:1000 in 3% BSA-TBS for 1 h at room temperature. Membranes were washed two times with TBST, followed by incubation with 1:2500 dilution of goat-anti mouse IgG (whole molecule)-Peroxidase conjugate (A4416, Sigma Aldrich) for 1 h at room temperature, followed by two TBST washes and developed with ECL Select™ WB detection reagent (Amersham™, Cytiva). Chemiluminescent exposure was performed simultaneously for three mouse sera-probed blots (i.e., nonimmunized and immunized) per experiment. Protein bands from corresponding blots were quantified using ImageJ by converting the original image to grayscale (8-bit) and measuring the mean gray value of the largest top-most protein band (i.e., between ∼30-180 kDa) with fixed measurement criteria. Background mean gray measurements were taken underneath the band located at 17 kDa with fixed measurement criteria. Net protein values were obtained by subtracting the background from the protein band mean gray value. Experiment performed in technical triplicate.

The murine antibody response produced from immunized mice was assessed for cross-reactivity to fungal pathogens, including a *C. neoformans* clinical isolate of strain H99 serotype A, *Aspergillus fumigatus* ATCC 204305, *Candida glabrata* BG87, and *C. albicans* SC3514. Protein was obtained from whole cell lysates by probe sonication in 100 mM Tris-HCl (pH 8.5) with a protease inhibitor cocktail tablet (Sigma-Aldrich) and 2% final SDS concentration. Proteins were quantified using the BCA method and were processed for western blot analysis as described above. Experiment completed in technical triplicate.

### CipC homology bioinformatic analyses

Basic Local alignment Search tool (BLAST)-p algorithm (https://blast.ncbi.nlm.nih.gov/Blast.cgi) was used to compare the protein sequence of CNAG_03007 of *C. neoformans* against the NCBI collection of non-redundant protein sequences (update date: 2024/04/20; number of sequences: 722063114). The search was performed with a 5000-search limit, the BLOSUM62 matrix scoring function, and an Expect threshold (e-value) of 0.05. Representative homologs selected from major fungal pathogen taxa with a >50% homologous identity for Clustal Omega multiple sequence alignment (MSA)^106^. Neighbour-joining tree phylogenetic analysis calculated using MEGA11, using Bootstrap testing and 1000 replications^107^.

## Author Contributions

J.G.-M. conceptualized the study. B.B. & J.G.-M. designed the study. B.B., N.C., H.W., A.S., D.C.L, & M.W. performed experiments. B.B., N.C., H.W., A.S., & D.C.L, contributed data to figures. B.B.& J.G.-M. performed data analysis. B.B. & J.G.-M. designed and developed final figures. B.B., & J.G.-M. wrote the manuscript. B.B. & J.G.-M. edited the manuscript. All authors have read and approved the submitted manuscript.

## Acknowledgments

Thank you to members of the Geddes-McAlister lab for their informative and constructive feedback on project design and manuscript preparation, as well as Samanta Pladwig, MSc. for Southern blot analysis and Wan Yun Zhu, BSc. for colony plate preparation and counting. We thank Dr. Cezar Khursigara (University of Guelph) for their technical assistance and use of microscopy instrumentation, Drs. John Perfect and Jennifer Tenor (Duke University) for the *C. neoformans* H99 clinical isolate, Dr. J.P. Xu (McMaster University) for the NAT resistance cassette, Dr. James Fraser (University of Queensland) for pSDMA58, Dr. Felix Meissner for providing the immortalized BALB/c macrophages, and Dr. Dyanne Brewer for operation of the mass spectrometer at the University of Guelph Advanced Analysis Centre – Mass Spectrometry Facility.

## Data Availability

The. RAW and affiliated files are deposited into the publicly available PRIDE partner database for the ProteomeXchange consortium.

PRIDE ID: PXD051506

Reviewer login

Password: OABkby1N

PRIDE ID: PXD051675

Reviewer login

Password: kiGW6jXG

## Supplemental Material

Fig. S1: Infectome profiling.

Fig. S2 Confirmation of macrophage-induced fungal protein abundance increases.

Fig. S3: *C. neoformans* infection-associated candidates display virulence deficits during *in vitro* phenotyping.

Fig. S4. Sequence alignment and Neighbor-joining tree of CipC with predicted homologs of fungal pathogens.

Fig. S5: Susceptibility of *cipC*Δ to antifungals.

Fig. S6: Co-immunoprecipitation of CipC and macrophage. Fig. S7: Loading control prior to sera exposure.

Table S1: Significantly different host proteins upon infection with *C. neoformans*.

Table S2. *C. neoformans* proteins identified from infectome profiling with exclusive detection or >16-fold increase in abundance.

Table S3: Proteins with significantly increased abundance upon deletion of cipC.

Table S4: Host proteins exclusively identified from CipC co-immunoprecipitation assay Table S5: Proteomes of extracellular vesicles derived from WT or cipC deletion strain

Table S6: Predicted antigenic proteins identified in extracellular vesicles derived from the cipC deletion strain

Table S7: Primers and Plasmids for gene deletion and complementation.

