## Supplementary material for "Systematic Infectome–Phenome Profiling Reveals Cryptococcal Infection-Associated Proteins Driving Immune System Remodeling and Immunization Potential": Supp. Figures

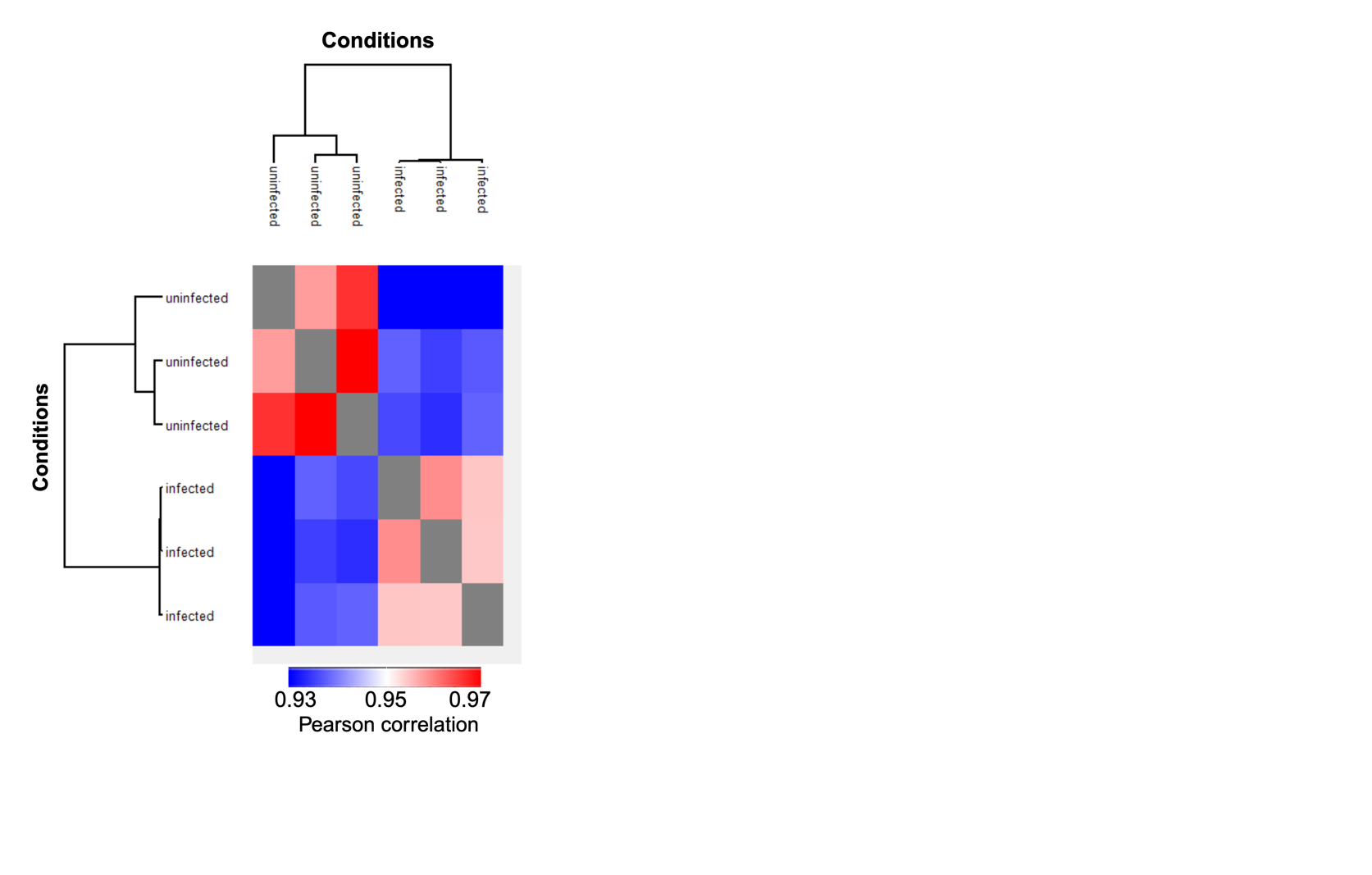

**Fig. S1: Infectome profiling.** Column correlation heat map of hierarchical clustering based on Euclidean distance of biological replicates for infected (i.e., *C. neoformans* co-cultured with macrophage) and uninfected (i.e., macrophage only) conditions. Experiment completed in biological triplicate.

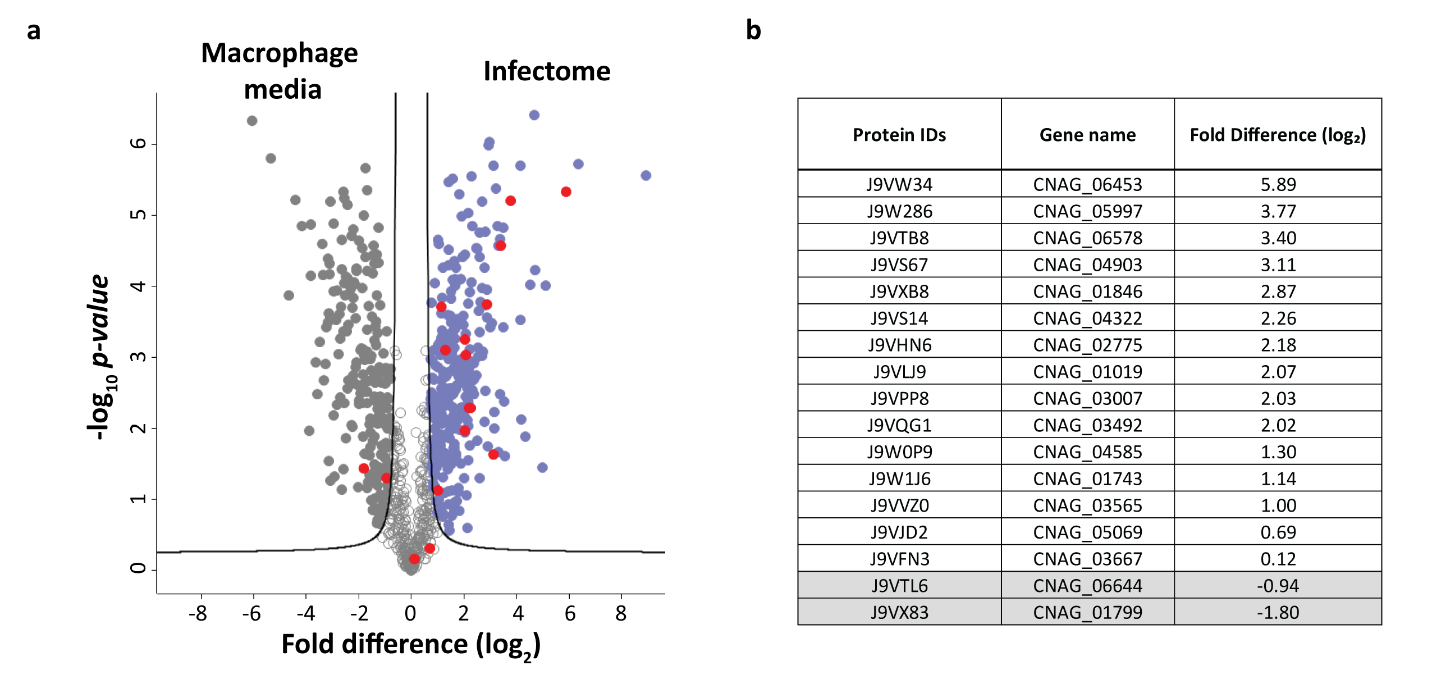

**Fig. S2: Confirmation of macrophage-induced fungal protein abundance increases. A)** Volcano plot comparison of *C. neoformans* in macrophage-infection mimicking conditions (i.e., 1.2 x 10^8^ fungal cells plated in tissue culture plates, 3 h incubation in DMEM, 37 °C, 5% CO_2_) and *C. neoformans* infection of BALB/c macrophages (i.e., infectome conditions; 1.2 x 10^8^ fungal cells co-cultured with 1.2 x 10^6^ macrophages for 3 h in DMEM, 37 °C, 5% CO_2_). Fungal proteins highlighted in red signify the 17 selected fungal proteins for phenotypic characterization. Student’s *t* test, *P* < 0.05; FDR = 0.05; S_0_ = 1. Experiment completed in biological quadruplicate. **B)** Log_2_ fold difference of 17 candidate fungal proteins from Infectome vs. Macrophage media volcano plot comparison. Proteins highlighted in gray indicate abundance profiles that are significantly higher in macrophage media conditions.

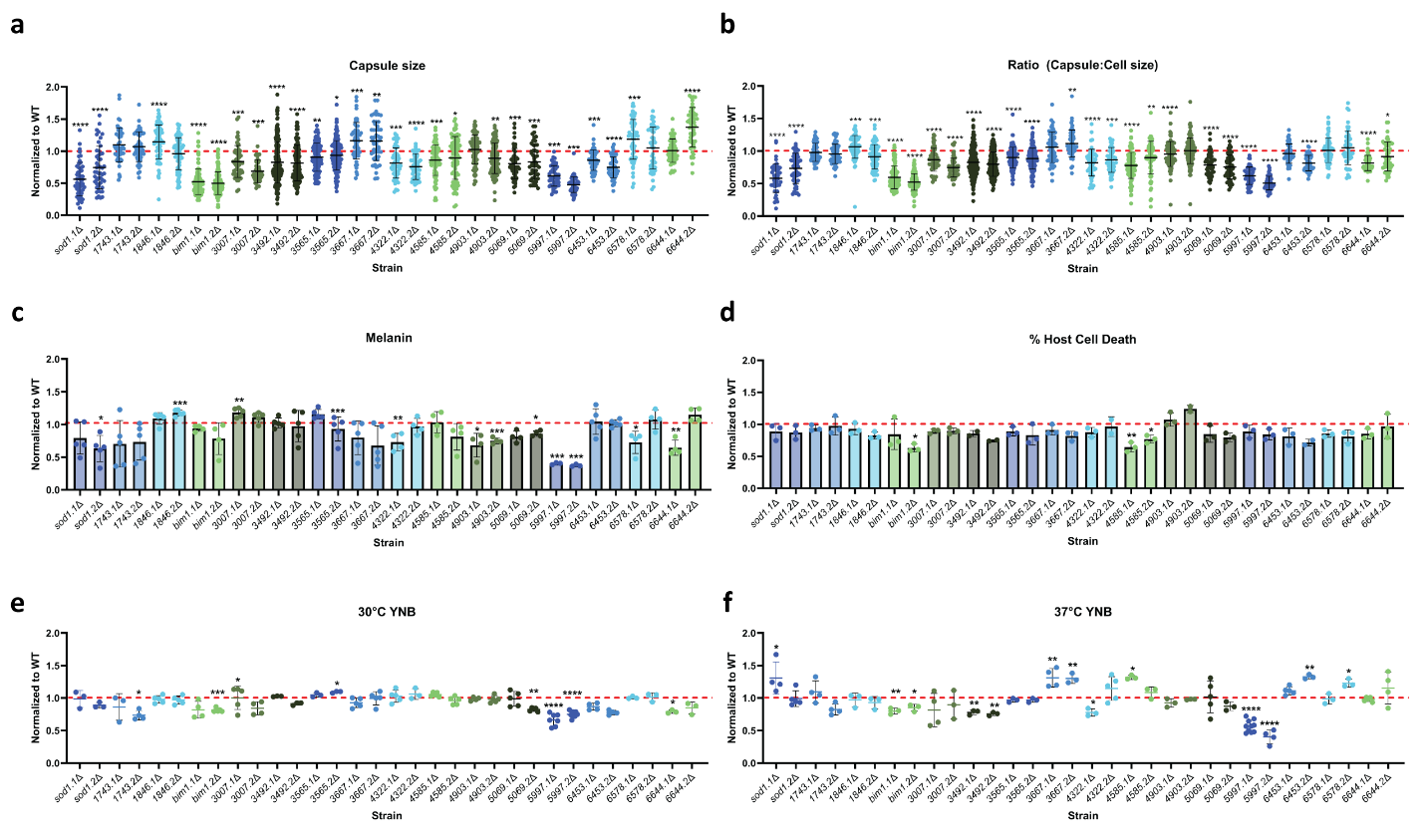
**Fig. S3: *C. neoformans* infection-associated candidates display virulence deficits during *in vitro* phenotyping. A)** Normalized capsule thickness of *C. neoformans* mutant strains against WT capsule size. Capsule assays performed in LIM following 24 h incubation. **B)** Normalized ratio between capsule thickness and cell size diameter for infection-associated candidate strains against WT. A minimum of 50 cells were measured for capsule and cell size assays across three biological replicates. **C)** Normalized percentage of melanin (pigment) produced by infection-associated strains against WT. Experiment performed in biological duplicate and technical triplicate. **D)** Percentage of host cell death produced following quantification of LDH release at 27 hpi of infection-associated mutant strains co-culture with BALB/c macrophages normalized against WT-induced host cell cytotoxicity. Experiment completed in biological triplicate and technical duplicate. Values shown are representative of one experiment. Normalized area under the curve (AUC) measurements from **(E)** 30 °C and **(F)** 37 °C growth profiles of mutant strains against WT following culture in YNB. Experiment completed in biological triplicate-quadruplicate and technical duplicate. Values shown are representative of one experiment. Statistical significance test between comparisons to WT by Student’s *t*-test: *P* < 0.05, *; *P* ≤ 0.01, **; *P* ≤ 0.005, ***; *P* ≤ 0.0001, ****.

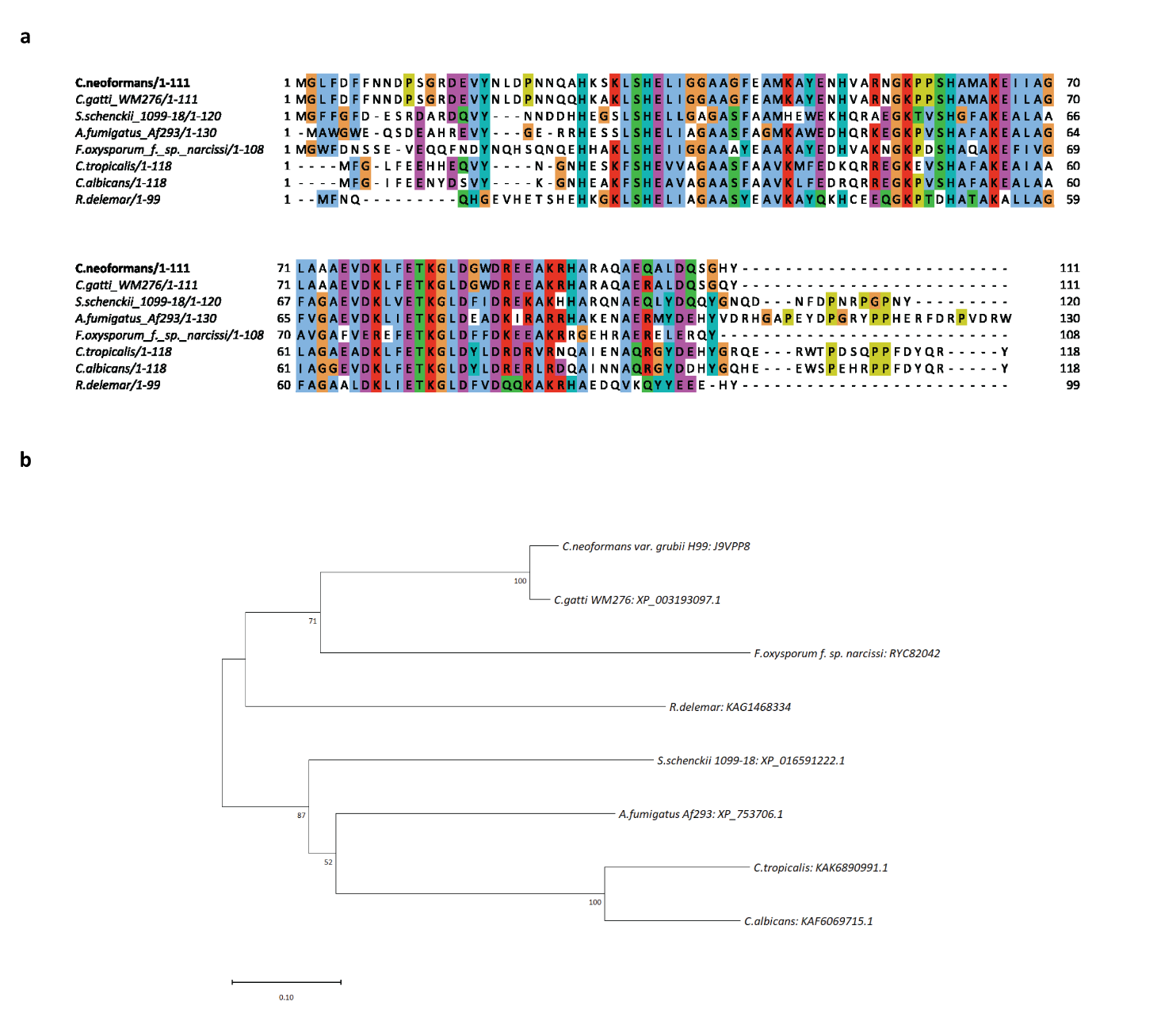

**Fig. S4. Sequence alignment and Neighbor-joining tree of CipC with predicted homologs of fungal pathogens. A)** CipC homologous amino acid sequences from *C. neoformans*, *Cryptococcus gatti* WM276, *Fusarium oxysporum f. sp. narcissi, Rhizopus delemar, Sporothrix schenckii, Aspergillus fumigatus* Af293, *Candida tropicaliss,* and *Candida albicans* were aligned using Clustal Omega software and visualized using Jalview. Amino acids coloured following Clustal X colour scheme: hydrophobic (blue), positive charge (red), negative charge (magenta), polar (green), cysteines (pink), glycines (orange), prolines (yellow), aromatic (cyan), unconserved (white). **B)** Evolutionary history of fungal pathogen CipC homologs inferred using the Neighbor-Joining method. Bootstrap values from 1000 replications are given at nodes. The scale bar represents 10% weighted sequence divergence. Evolutionary distances were computed using the Poisson correction method and are in the units of the number of amino acid substitutions per site, involving 8 amino acid sequences. All ambiguous positions were removed for each sequence pair (pairwise deletion option). Analysis conducted in MEGA11.

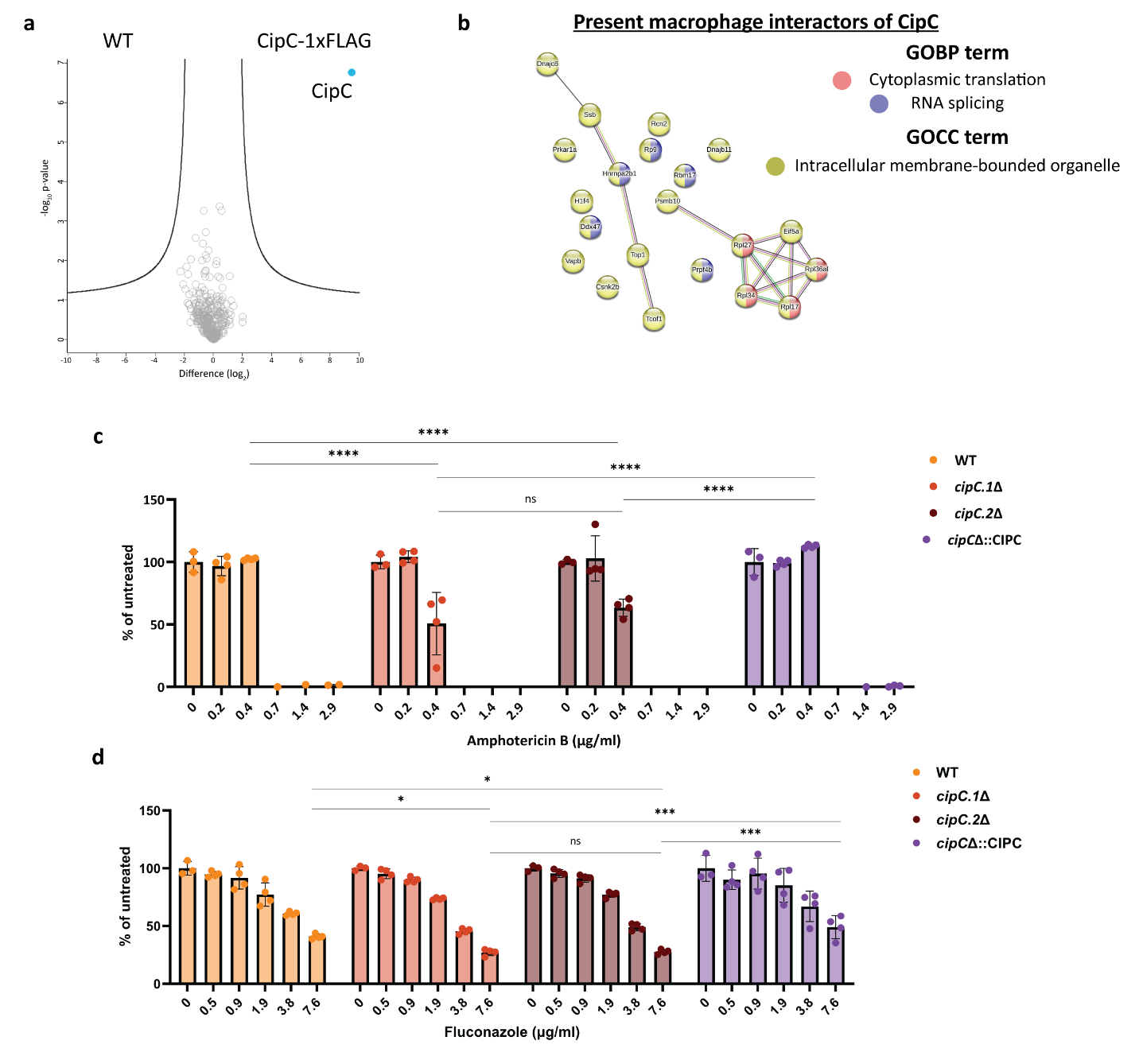

**Fig. S5: Susceptibility of *cipC*Δ to antifungals.** Microdilution assay of amphotericin B sensitivity to strains at concentrations of 0-2.9 µg/mL and fluconazole sensitivity to strains at concentrations of 0-7.6 µg/mL. Cell density was measured at OD_600nm_ and represented as percentage of growth of the untreated control. Data shown representative of one experiment. Experiment completed in biological quadruplicate and technical duplicate. Statistical significance assessed with Tukey’s multiple comparisons test: ns = not significant; *P* < 0.05, *; *P* ≤ 0.0005, ***; *P* ≤ 0.0001, ****.

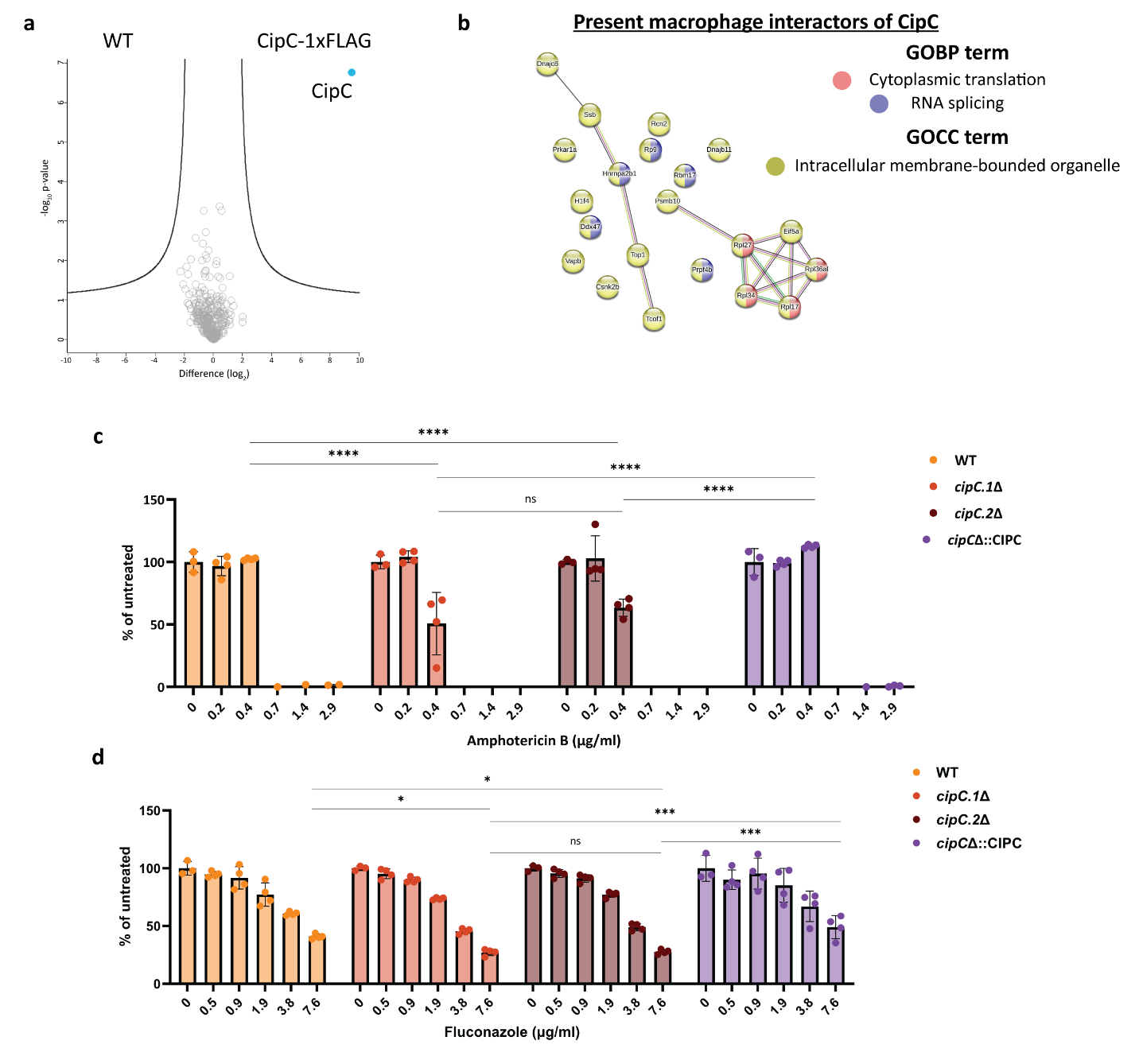

**Fig. S6: Co-immunoprecipitation of CipC and macrophage.** Interactome volcano plot of *cipC*Δ::CIPC-1x FLAG at 3 hpi with macrophage (MOI 100) versus WT *C. neoformans* (negative control). Grey (host) and blue (fungal) proteins highlighted. Statistical analysis of volcano plot by Student’s *t*-test (*p*-value ≤ 0.05; FDR = 0.05; S_0_ = 1). Co-immunoprecipitation experiment completed in biological quadruplicate.

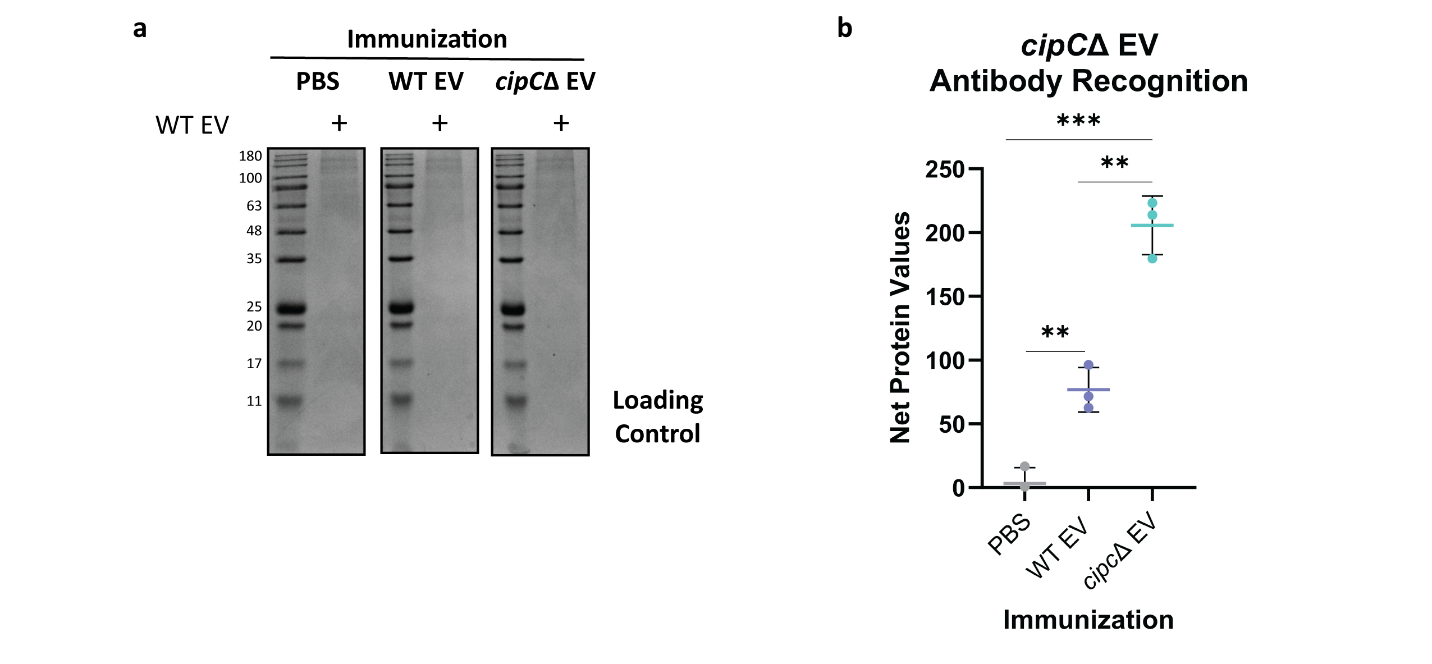

**Fig. S7: Loading control prior to sera exposure.** Western blot of WT-derived EVs. Equal EV protein content indicated as loading control. Experiment completed in technical triplicate.
